# Central amygdala Isl1 neurons control biting by integrating sensory and motivational signals

**DOI:** 10.64898/2026.02.03.703447

**Authors:** Wenyu Ding, Alisson Pinto de Almeida, Wenkang Wang, Chunyu Qu, Rüdiger Klein

## Abstract

Efficient orofacial motor control is essential for adaptive interactions with the environment, yet the neural substrates that regulate the force and precision of biting remain poorly understood. Here, we identify a subpopulation of neurons in the central amygdala (CeA) expressing the transcription factor *Isl1* (CeA^Isl1^) that plays a crucial role in modulating biting behavior in mice. In vivo calcium imaging revealed that CeA^Isl1^ neurons are robustly activated at the onset of biting across materials of varying physical properties, with distinct neuronal ensembles selectively encoding responses to the physical properties of the objects. CeA^Isl1^ neuronal activity scales positively with the hardness of the object, suggesting a role in force modulation. Optogenetic activation of CeA^Isl1^ neurons enhances biting behavior toward edible or non-edible objects, induces fictive feeding in the absence of physical targets and exerts a reinforcing effect on behavior, whereas inhibition of CeA^Isl1^ neurons impaired efficient biting of solid food by reducing jaw-closing muscle activity. Projection-specific manipulations revealed that activation of CeA^Isl1^ projections to the parvocellular reticular formation (PCRt) and pedunculopontine tegmental nucleus (PPtg) increased the duration and frequency of biting, with CeA^Isl1^-to-PPTg stimulation producing a positive motivational valence. These findings uncover a previously unrecognized sensorimotor function of the central amygdala in calibrating bite force and precision, linking motivational states to skilled motor output.

## Introduction

Feeding behavior is controlled by many brain circuits and neuron types^1-8^. This is because the brain has to integrate the many sensory and physical attributes of the food with the interoceptive inputs from the body. The central amygdala (CeA) has emerged as a brain hub regulating feeding behavior^8-13^. The CeA is composed of distinct microcircuits that evaluate appetitive stimuli to drive reward and consummatory behaviors^8,9^, but also separate microcircuits that suppress appetite when reaching satiety and in the presence of nausea^10,11^. The CeA has also been shown to drive hunting-like behavior, including pursuit, grabbing and biting^14^. However, it is unclear if in addition to the known appetitive microcircuits, the CeA contains distinct biting neurons. Mapping the CeA cell types and microcircuits that respond to the variety of sensory attributes (taste, smell) and physical properties (texture, density, viscosity) of food and process this information to generate the appropriate motor output is important for understanding how consumption of solid food is regulated.

Feeding requires precise coordination of jaw movements. These movements must adapt to different food textures and densities. Disturbances in their control are linked to eating disorders. Understandig how the brain regulates jaw movements is therefore critical for both basic neuroscience and clinical applications. The tongue/jaw-related primary motor cortex (tjM1) plays a central role in initiating and controlling jaw movements and biting force^15-19^. Distinct excitatory neuron populations within tjM1 contribute to the generation of specific jaw motor patterns^20,21^. Subcortical structures also shape orofacial behaviors. The CeA has long been known to evoke jaw opening, biting, and chewing across species ^22-24^. Yet, the identity of the specific cell types involved remained unclear.

Previous studies demonstrated that CeA biting neurons project to the parvocellular reticular formation (PCRt). This region contains mandibular and cervical premotor neurons that mediate biting attacks^14^. More recent work showed that CeA projections to the parabrachial and supratrigeminal nuclei elicit rapid orofacial behaviors that drive ingestion of both food and non-food objects^25^. Together, these findings point to the CeA as a key hub for orchestrating orofacial actions.

We hypothesized that the CeA not only initiates jaw movements but also modulates their intensity in response to sensory input. To investigate this possibility, we focused on identifying the neuronal population involved in jaw movements. The CeA is subdivided into three spatial subdivisions, the capsular (CeC), lateral (CeL) and medial subdivisions (CeM). The CeM is the major output region and it has been implicated in attack-related biting through its projections to the PCRt^14^. The so far best characterized appetitive CeM population marked by expression of the serotonin receptor Htr2a (and partially overlapping with the Pnoc population) promotes palatable food consumption and positive reinforcement^8,9,12,13^. But unlike the biting neurons, CeM^Htr2a^ neurons mediate their functions via the parabrachial nucleus (PBN) and were therefore an unlikely candidate for the biting neurons.

Recent single-cell RNA sequencing studies have highlighted a CeM population marked by the LIM-homeodomain transcription factor *Isl1*, having little overlap with CeM^Htr2a^ or Pnoc neurons^26^. Isl1 neurons were previously described as a major CeM population in developmental tracing studies^27^ and were shown to project to multiple hindbrain target regions, including PCRt^26,28^. However, their function in the regulation of feeding or biting behavior remained unknown.

Here, we used Isl1-CreER mice to selectively target this genetically defined CeM population and examined its role in modulating biting behavior. We found that CeA^Isl1^ neurons are robustly activated at the onset of biting and encode the physical properties of food, with harder objects eliciting stronger population responses. Single-neuron analysis revealed functionally distinct subpopulations tuned to material-specific sensorimotor features. Activation of CeA^Isl1^ neurons increased the vigor of orofacial movements without altering food intake, whereas their inhibition impaired efficient biting of solid food. Projection-specific manipulations demonstrated that activation of CeA → PCRt and CeA → pedunculopontine tegmental nucleus (PPTg) pathways increased the duration and frequency of biting behavior. Notably, CeA → PPTg stimulation enhanced biting with positive motivational valence, identifying a parallel pathway that reinforces orofacial actions through emotional drive.

## Results

### CeA neurons show robust activation at bite onset across diverse objects

Using the known localization of CeA^Pkcδ^ neurons to the CeL^8^, we confirmed that Isl1 immunoreactive neurons were largely confined to the CeM (Fig. 1A). We then used Isl1-CreER mice in combination with Isl1 immunostaining to show high fidelity of Cre expression, with strong overlap with Isl1 immunoreactivity (fig. S1, A to C). A comparison with other transcriptionally defined cell clusters revealed that the *Isl1*^+^ population was distinct from other CeA neuronal subtypes, including the *Sst* and *Pnoc* clusters previously associated with appetitive behavior ^9,12,29^ (fig. S1D).

**Figure 1.**
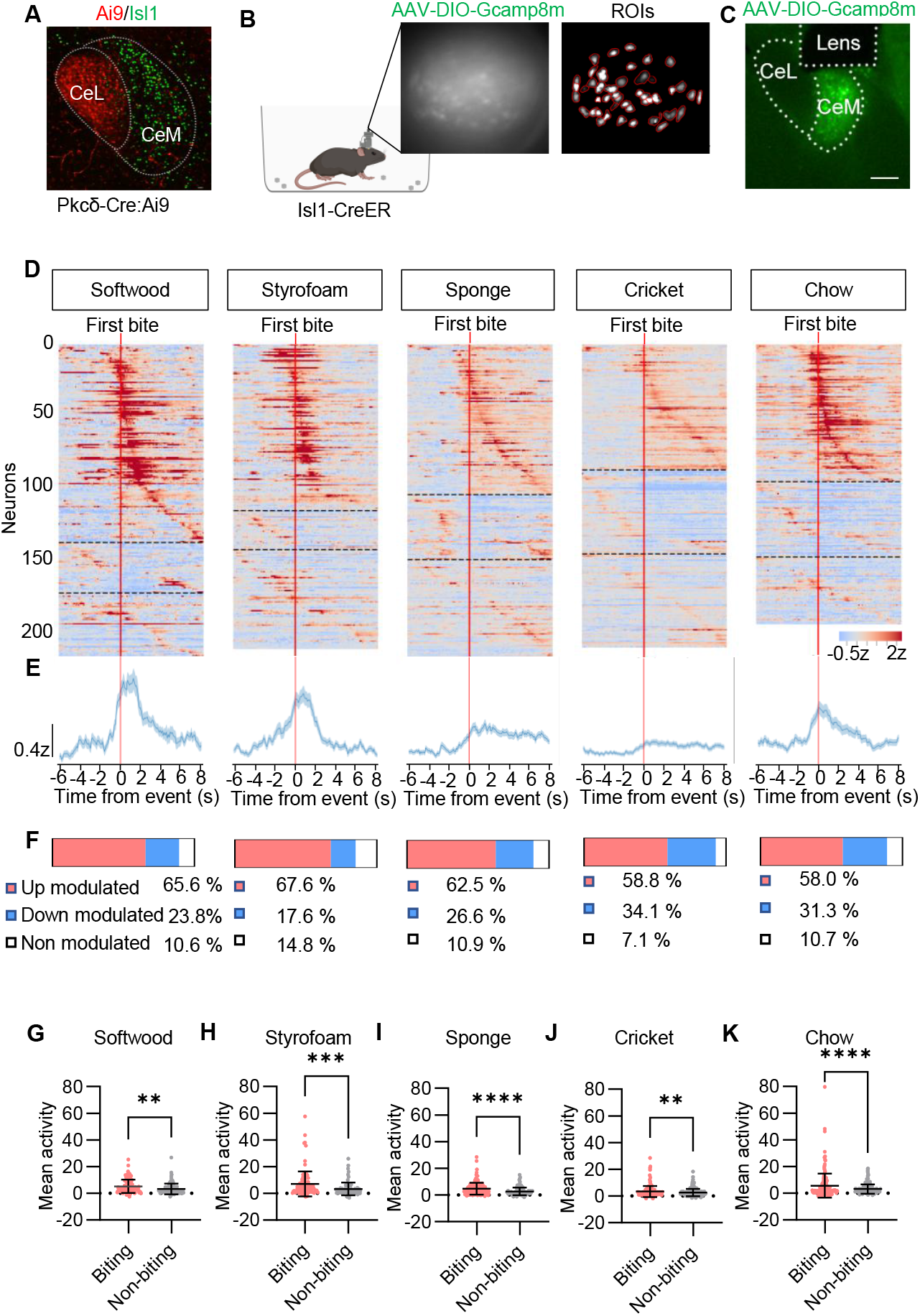
CeA^Isl1^ neurons responses during object-biting behaviors. **A**. Representative image showing Isl1 immunostaining in the CeA of Pkcδ-CreER : Ai9 mouse. Scale bar, 30 µm. **B**. Schematic of a freely moving mouse expressing GCaMP8m in CeA^Isl1^ neurons and carrying a head-mounted miniscope. Left panel: raw calcium imaging frame; right panel: manually segmented regions of interest (ROIs) corresponding to individual neurons. **C**. Post hoc histological validation showing GCaMP8m expression (green) in the medial central amygdala (CeM), with the lens track visible above the CeA. Scale bar, 100 µm. **D**. Heatmaps of individual neurons (rows) showing z-scored calcium activity (color scale) aligned to the first bite (vertical red line) of each substrate: softwood, Styrofoam, sponge, cricket, and chow. Neurons are sorted by response type. N= 12 mice per group. **E**. Average responses of the z-scored calcium activity for each condition. Shaded areas represent s.e.m. **F**. Proportions of neurons classified as up-modulated (red), down-modulated (blue), or non-modulated (white) for each substrate. **G–K**. Mean neuronal activity during object-biting versus non-biting epochs: soft wood (**G**), Styrofoam (**H**), sponge (**I**), cricket (**J**), and chow (**K**) (n= 12 mice per group. Wilcoxon signed-rank test; **P ⍰ <10.01, ***P ⍰ <10.001, ****P ⍰ <10.0001).

To assess whether the activities of individual CeA^Isl1^ neurons were associated with biting, we performed *in vivo* calcium imaging in freely behaving mice. We delivered Cre-dependent GCaMP8m-expressing virus unilaterally into the CeA of Isl1-CreER mice, followed by implantation of a gradient index (GRIN) lens in the same location. Calcium signals in the infected neurons were recorded using a head-mounted miniscope (Fig. 1, B and C). We tested animals that were fasted for four hours, under five separate conditions: exposure to non-food items such as softwood, Styrofoam, sponge, as well as live crickets, and normal chow. Recordings were done for 5 min each. To analyze how CeA^Isl1^ neurons responded at the onset of biting, we aligned calcium activity traces to the “first bite”, defined as the initial bite of each discrete bout of biting behavior, across the five conditions (Fig. 1, D and E). Notably, although the population mean activity revealed a prominent peak at bite onset, the response profiles differed between materials, with harder substrates (softwood, Styrofoam, chow) evoking sharper, higher amplitude activation compared to softer substrates (sponge, cricket), regardless of whether they were edible or inedible (Fig. 1E). In all cases, a large proportion of neurons showed clear modulation around bite onset, with consistent recruitment patterns across both edible and inedible substrates. Up-modulated neurons comprised the majority of the population in each condition (ranging from 58.0 % in chow to 67.6 % in Styrofoam), while smaller subsets were down-modulated or non-responsive (Fig. 1F). While the frequency of bite events did not differ across materials (fig. S1E), the duration of individual bites varied substantially, with softer objects eliciting longer bite durations (fig. S1F). This dissociation raised the possibility that the observed neural responses were not a reflection of how much the animal preferred or enjoyed biting a particular object. Instead, the activity more likely reflected biomechanical demands or motor output features such as bite force or muscle engagement associated with different material properties. For example, soft wood was bitten for less time due to high resistance (fig. S1F), but the population mean activity was higher than for the others (Fig. S1G). The temporal dynamics of CeA^Isl1^ activation (peaking within seconds after the bite) were preserved across conditions. These findings indicate that CeA^Isl1^ neuron activity was closely tied to the execution of biting behavior and may encode material resistance during biting, as activity was consistently elevated during biting compared to non-biting periods (Fig. 1, G to K).

To examine how neuronal activity related to feeding phases, we analyzed the structure of chow consumption. Mice typically began with gnawing, forceful oral interactions near the mouth, and in ∼1-11 % of the time this was followed by rhythmic chewing after moving the pellet outward (fig. S1H). Population mean activity was significantly higher during gnawing than during chewing or non-gnawing periods (fig. S1, I and J), with no difference between chewing and non-chewing epochs (fig. S1K). Correspondingly, 47.4 % of neurons were modulated during gnawing, but only 2.9 % during chewing (fig. S1, L and M). Thus, CeA^Isl1^ activity was preferentially engaged during the initial interaction with food, particularly first bite and gnawing, and largely unresponsive to subsequent chewing, suggesting a role in encoding the sensory–motor demands of bite initiation rather than general food processing.

To assess CeA^Isl1^ neuron activity during ethologically relevant orofacial behaviors, we performed calcium imaging in freely moving mice during cricket hunting. We defined pursuit as the period during which the mouse actively followed the cricket prior to capture, and holding as the epoch immediately following capture, when the mouse restrained the cricket in its mouth and often relocated to a secure area before consumption. We found that the average activity across the population did not differ significantly between these behavioral epochs (fig. S1, N and O), likely because all three involved orofacial engagement to some extent. Pursuit often included attempted strikes, and holding required continued mouth contact with the prey. A significantly higher proportion of neurons was modulated during gnawing (fig. S1P to R), suggesting that CeA neurons were more consistently engaged during forceful orofacial actions than during pursuit or prey holding.

### Distinct populations of CeA neurons encode material-specific biting behavior

Next, we asked whether distinct subpopulations of CeA^Isl1^ neurons exhibited material-specific activity patterns reflecting the physical properties of the bitten objects. We performed longitudinal cell registration across conditions for the previously shown data. Heatmaps of single-neuron calcium activity revealed diverse response profiles depending on the object the mice interacted with. Neurons were grouped into five functional clusters based on their dominant activation near the onset of the first bite. Strikingly, these clusters exhibited distinct material preferences: Cluster 1 responded most strongly to silicone, Cluster 2 was preferentially activated by softwood, Cluster 3 responded to both sponge and Styrofoam, but showed greater activity for sponge, and Cluster 4 was selectively activated by Styrofoam. Cluster 5 showed non-responsive cells. These patterns were consistent across animals and stable across trials (Fig. 2A).

**Figure 2.**
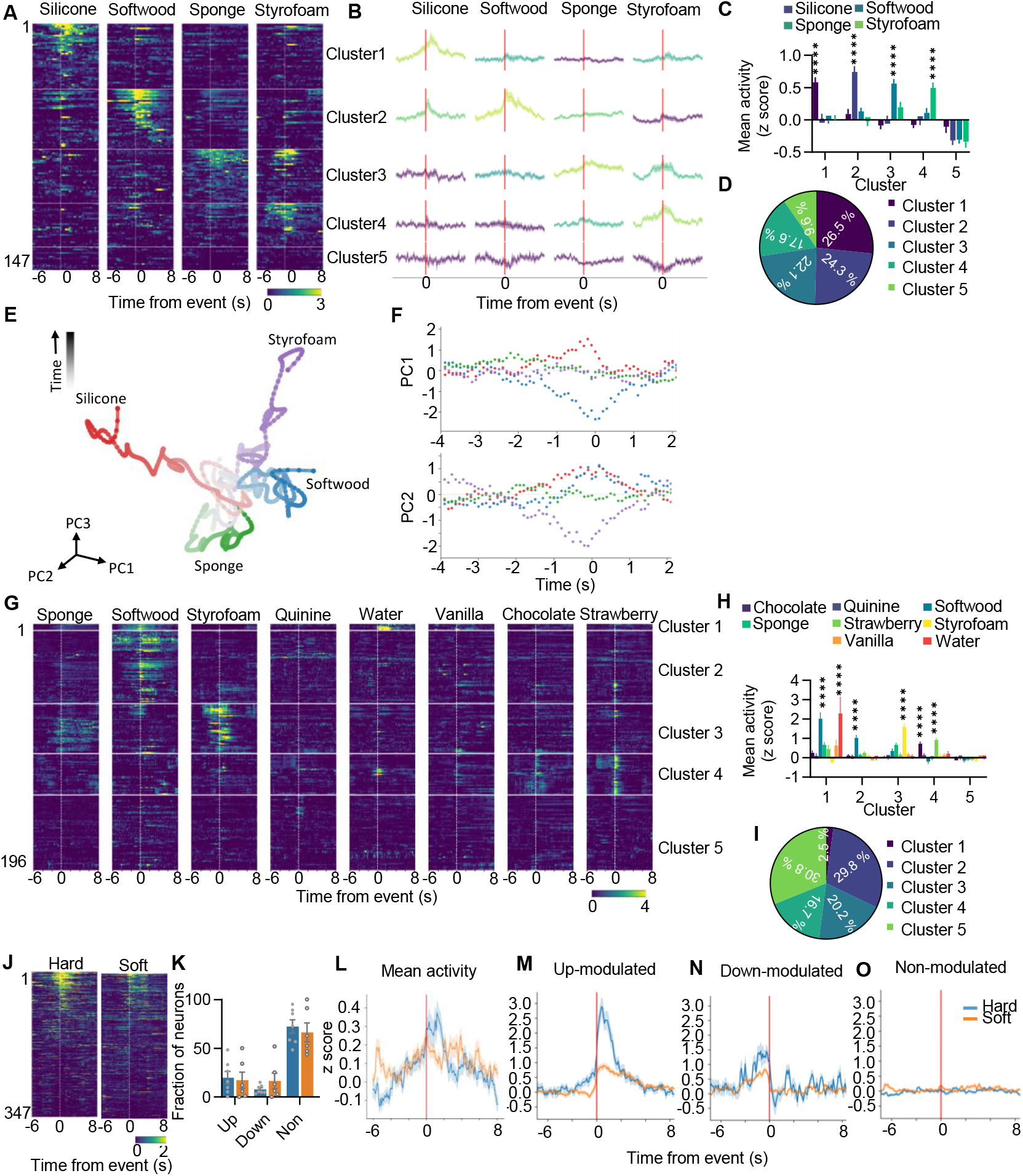
The material-specific encoding properties of CeA ^Isl1^ neurons. **A**. Heatmaps of the response to different materials. Each row represents the activities of one neuron. Neurons were grouped by clustering based on their mean response profiles across conditions (n= 6 mice). **B**. Average activity traces for each cluster. Distinct temporal response patterns and stimulus selectivity emerged across clusters. Shaded areas represent s.e.m. **C**. Neuronal activity across five functional clusters during different bite conditions: silicone, softwood, sponge, and Styrofoam. (n= 6 mice per group; ****p < 0.0001; two-way ANOVA with Tukey’s post hoc test). **D**. Proportion of neurons in each cluster. **E**. Principal component analysis (PCA) of trial-averaged population activity across mice reveals distinct trajectories for bites on silicone (red), sponge (green), softwood (blue), and Styrofoam (purple). Each trajectory represents the temporal evolution of population activity projected onto the first three principal components (PC1–PC3), with time progressing along the trajectory (arrow). **F**. Time courses of the first two principal components (PC1 and PC2) aligned to bite onset. Distinct material-specific dynamics emerge around the time of biting, indicating that CeA^Isl1^ ensemble activity transiently encodes the physical identity of the bitten material. **G**. Heatmaps of the response to different stimuli. Each row represents the activities of one neuron. Sponge, softwood, and Styrofoam aligned to bite onset. Quinine, water, vanilla milk, chocolate milk and strawberry milk aligned to lick onset. Neurons were grouped by clustering based on their mean response profiles across conditions (n= 6 mice). **H**. Neuronal activity across five functional clusters during different conditions: sponge, softwood, Styrofoam, quinine, water, vanilla milk, chocolate milk and strawberry milk. (n= 6 mice per group, ****p < 0.0001, two-way ANOVA with Tukey’s post hoc test). **I**. Proportion of neurons in each cluster. **J**. Heatmaps of responses to hard silicone and soft silicone. Each row represents the activities of one neuron (n = 6 mice). **K**. Fraction of neurons classified as up-modulated, down-modulated, or non-modulated. Blue is hard silicone, orange is soft silicone (n= 6 mice; N.S.; two-way ANOVA with Tukey’s post hoc test). **L-O**. Average responses of all neurons (**L**) and neurons classified as up-modulated (**M**), down-modulated (**N**), or non-modulated (**O**). Shaded areas represent s.e.m.

Cluster-averaged activity traces confirmed that responses were time-locked to bite onset and differed in both amplitude and dynamics depending on the material (Fig. 2B, fig. S2, A and B). Functional clustering of CeA^Isl1^ neurons revealed five distinct response profiles across bite conditions (Fig. 2C), with neurons distributed across clusters in a largely balanced manner, except for a smaller non-responsive cluster (Fig. 2D). The spatial arrangement of these clusters within the CeA^Isl1^ population appeared random (fig. S2C). Together, these results suggest that CeA^Isl1^ neurons contain functionally distinct subpopulations that encode material-specific sensory-motor features, supporting a role for the central amygdala in evaluating the physical properties of objects during orofacial behavior.

To examine how CeA^Isl1^ population dynamics encoded the physical properties of different materials, we performed principal component analysis (PCA) on the pooled, trial-averaged calcium activities across all recorded neurons and mice. The resulting low-dimensional trajectories revealed distinct population activity patterns associated with biting different materials (Fig. 2E). When projected onto the first three principal components (PC1–PC3), trajectories corresponding to silicone, sponge, softwood, and Styrofoam diverged in state space around the time of biting, indicating material-specific neural representations. Following the bite, trajectories gradually converged, suggesting that CeA^Isl1^ population activity transiently differentiated object identity during the biting action before returning toward a common post-event state (Fig. 2F).

To test whether neurons differentiated between flavors during food consumption, we analyzed calcium activity of CeA^Isl1^ neurons aligned to bites of food pellets immersed in strawberry, vanilla, or chocolate aroma, water, or quinine. The same neuronal population was tracked across all conditions, and neurons were clustered based on their response dynamics relative to bite onset (fig. S2D). Most responsive neurons exhibited highly similar activation patterns regardless of flavor, with little evidence for selective responses to specific sensory identities (fig. S2, E and F). Similar results were obtained with non-edible sponges infused with the same flavors (fig. S2, G to J). This response pattern was consistent even for quinine, suggesting that CeA^Isl1^ neurons were not tuned to flavor identity but instead encoded general features of food-contact behavior, such as sensory detection or motor engagement independent of the specific taste or aroma. Frequency and duration of biting behavior did not differ significantly across flavors for either food or sponges (fig. S2, J to M), suggesting that the observed neural responses were not related to changes in motor output or behavioral preference.

Licking and biting are both fundamental motor actions required for consummatory behavior and previous studies have shown that neurons in the CeA respond to licking^12,30,31^. To determine whether CeA^Isl1^ neurons responded to different liquids, we performed calcium imaging while mice were presented with flavored or unflavored liquids (water, quinine, chocolate milk, vanilla milk, strawberry milk). We registered individual neurons across all conditions including different hard objects (sponge, softwood, Styrofoam) to enable within-neuron comparisons and clustered activity traces by response dynamics (Fig. 2G). We identified five distinct neuronal clusters with event-aligned activity profiles across the different substrates. A small cluster 1 exhibited an intermixed response pattern, with most neurons responding robustly to water and some to softwood. Since the mice were water-deprived for four hours, water likely acted as a rewarding stimulus in this context, though cluster 1 was distinct from the flavored milks. Cluster 2 was strongly activated by softwood, while Cluster 3 responded preferentially to sponge and Styrofoam. Cluster 4 neurons were selectively engaged during licking of all palatable liquids regardless of flavor (Fig. 2, H and I; fig. S3, A and B). No specific cluster was preferentially tuned to quinine, and all liquids evoked significantly higher activities during licking compared to non-licking periods (fig. S3, C to G). This suggests that the act of licking itself contributed to CeA^Isl1^ activation, even for less palatable or aversive liquids. Licking metrics revealed no significant differences in lick frequency or duration across liquid types (fig. S3, H and J). Together, these findings suggest that CeA^Isl1^ neurons include mostly material-specific and some reward-sensitive subpopulations, while also integrating motor-related signals. Furthermore, the partial overlap between softwood-responsive and water-responsive neurons suggests that some CeA^Isl1^ neurons may integrate sensory and hedonic cues during consummatory behavior.

### The activity of CeA neurons is positively correlated with the hardness of the bitten object

To examine if CeA^Isl1^ neural responses were correlated with biting force, we compared the calcium activities of CeA^Isl1^ neurons when the mice were biting the same object (silicone), but of different firmness. Population heatmaps revealed that a fraction of neurons was activated more strongly when biting hard compared to soft silicone (Fig. 2, J and K), consistent with more robust engagement or sensory feedback. Hard silicone elicited higher mean activity across the entire population and greater peak responses of up-modulated neurons than soft silicone (Fig. 2, L and M). Conversely, hard silicone elicited more pronounced suppression of down-modulated neurons relative to baseline than soft silicone (Fig. 2N). Non-modulated neurons remained flat across conditions (Fig. 2O). These results suggest that harder materials drive stronger and more diverse neural responses, potentially reflecting greater sensory input, increased biting force, or longer engagement durations. This finding supports the interpretation that CeA^Isl1^ neurons encode the mechanosensory features of biting in a material-specific manner, integrating both sensory feedback and stimulus identity.

### CeA neurons are sufficient to increase bite force and frequency

Given that CeA^Isl1^ neuronal activity correlated with the hardness of the bitten object, we hypothesized that these neurons may also scale with bite force. If so, artificial activation may enhance bite strength compared to no or weak stimulation. To test this, we optogenetically activated these neurons in Isl1-CreER mice bilaterally injected with a Cre-dependent AAV expressing ChR2 in the CeA and induced Cre expression by Tamoxifen (Fig. 3A). Mice were 4-hour food deprived and then presented with three 2-cm linguine pieces (a ribbon-shaped thick and relatively hard pasta that requires substantial bite force to break) per session, with or without 473 nm light stimulation across testing days (Fig. 3B). Optogenetic stimulation of CeA^Isl1^ neurons significantly increased the frequency of biting bouts compared to non-stimulated sessions, but only at 20 Hz (Fig. 3C), suggesting a frequency-dependent effect on motor behavior. Control mice expressing a Cre-dependent EGFP virus did not show this effect. Despite the increased bout number, the duration of individual feeding bouts was significantly shorter in ChR2-expressing mice compared to EGFP controls during 20 Hz light stimulation (Fig. 3D), suggesting less controlled or more forceful biting. Consistent with this, the total amount of linguine consumption was similar between groups (Fig. 3E), but ChR2 mice left more fragmented pieces scattered on the floor, suggesting that bites may have been overly forceful or imprecise. These results suggest that CeA^Isl1^ activation enhances the vigor of orofacial movements but may disrupt fine control of bite force, leading to inefficient food engagement without increased consumption.

**Figure 3.**
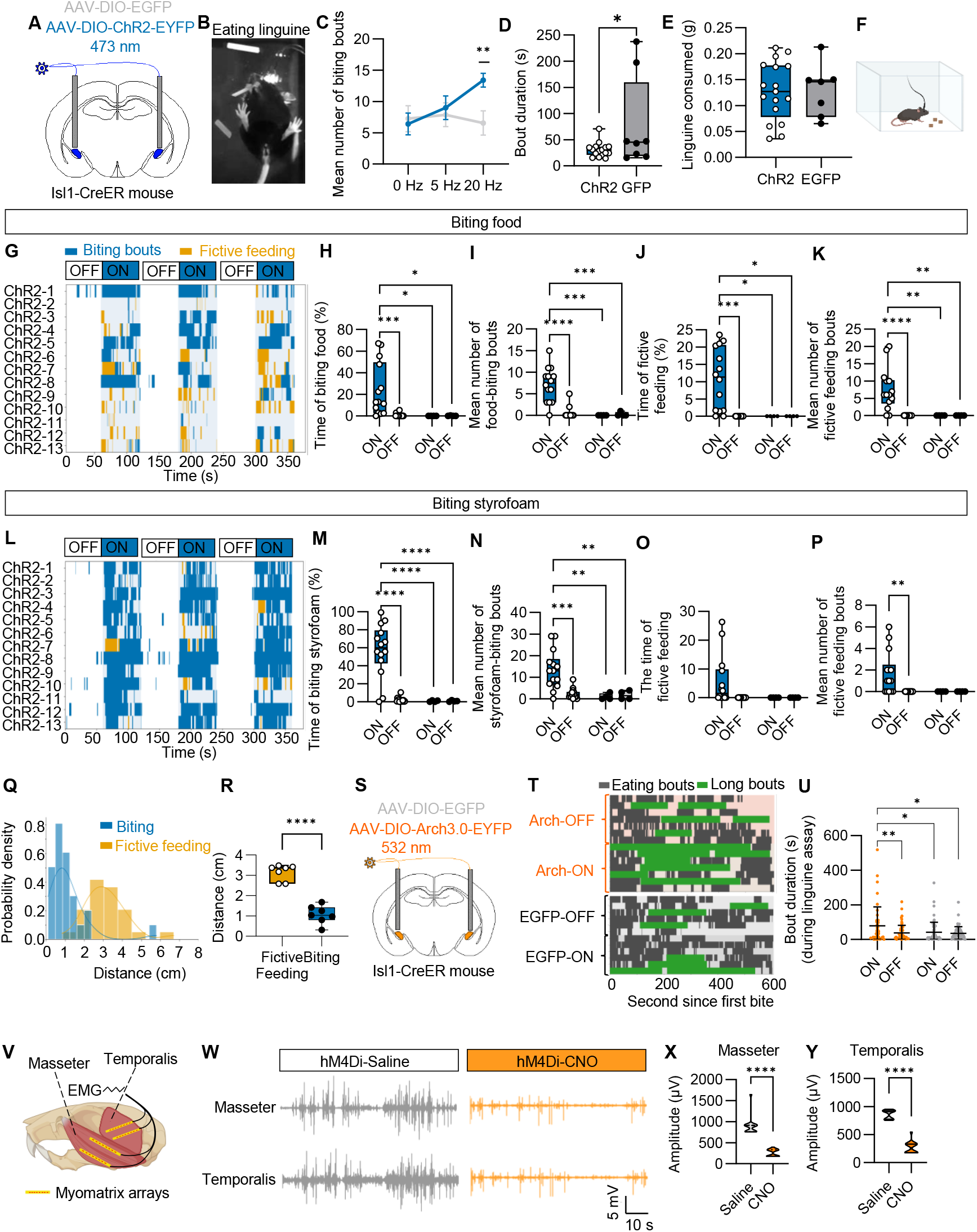
CeA ^Isl1^ neuron activation enhances biting efficiency, frequency, and fictive feeding behaviors. **A**. Schematic of optic-fiber placement above CeA^Isl1^ neurons expressing either ChR2 or EGFP. **B**. Representative frame from video showing a mouse grasping and biting linguine. **C**. Mean number of biting bouts across stimulation conditions (n= 12 mice per group; **p < 0.01, two-way ANOVA with Tukey’s post hoc test). **D**. Biting bout duration during 20 Hz stimulation (*p < 0.05, unpaired t-test). **E**. Weight of linguine consumed by 4 h fasted animals (N.S.; unpaired t-test). **F**. Schematic of a mouse freely biting pieces of chow/Styrofoam. **G**. Raster plot showing biting bouts (blue) and fictive feeding (orange) across individual fed ChR2 mice during alternating light OFF and light ON epochs in biting food assay. **H-K**. Quantification of behaviors during ON and OFF periods for both ChR2 mice and EGFP mice in biting food assay. **H**. Percentage of time spent biting food. **I**. Mean number of food-biting bouts. **J**. Percentage of time engaged in fictive feeding. **K**. Mean number of fictive feeding bouts. N= 13 mice per group. Two-way ANOVA with Tukey’s post hoc test, **\***P1<10.05, **P1<10.01, ***P1<10.001, ****P1<10.0001. **L**. Same as in panel **G** but for Styrofoam pieces. **M-P**. Quantification of behaviors during ON and OFF periods for both ChR2 mice and EGFP mice in biting Styrofoam assay. **M**. Percentage of time spent biting Styrofoam. **N**. Mean number of Styrofoam-biting bouts. **O**. Percentage of time engaged in fictive feeding. **P**. Mean number of fictive feeding bouts. N= 13 mice per group. Two-way ANOVA with Tukey’s post hoc test, **\***P ⍰ <10.05, **P ⍰ <10.01, ***P ⍰ <10.001,****P ⍰ <10.0001. **Q**. Probability density distributions of distances of the animal to the food source during biting (blue) and fictive feeding (orange) events. **R**. Quantification of mean distances for each behavior across animals (n = 7 mice, ****p < 0.0001, two-tailed unpaired t-test). **S**. Schematic of optic-fiber placement above CeA^Isl1^ neurons expressing either Arch3.0 or EGFP. **T**. Raster plots showing eating bouts (grey for eating bouts and green for eating bouts longer than 130 s) across light OFF and ON epochs in Arch3.0and EGFP groups during linguine consumption assays. **U**. Duration of individual feeding bouts across light OFF and ON epochs in Arch3.0 and EGFP groups during linguine consumption assays (n = 7 mice per group; \***p** < 0.05, **p < 0.01; two-way ANOVA with Tukey’s post hoc test). **V**. Schematic of Myomatrix microarrays (bottom), each containing eight electrode contacts on a flexible substrate, implanted into the masseter and temporalis muscles for electromyographic (EMG) recordings. **W**. Representative EMG traces from the right masseter and temporalis muscles of a freely behaving Isl1-CreER mouse expressing the inhibitory DREADD hM4Di in the central amygdala. Traces show activity during food biting after intraperitoneal injection of saline or CNO (2 mg kg^−1^). **X**. EMG amplitude measurements from the masseter muscle during food biting after saline or CNO (0.4 mg kg^−1^) injection in Isl1-CreER; hM4Di mice. n=4; ****p < 0.0001. **Y**. EMG amplitude measurements from the temporalis muscle during food biting after saline or CNO (0.4 mg kg^−1^) injection in Isl1-CreER; hM4Di mice. n=4; ****p < 0.0001.

To further determine whether CeA^Isl1^ neurons were sufficient to drive biting behavior, we presented fed mice with food or Styrofoam blocks (Fig. 3F). Blue light (473 nm) was delivered in an alternating epoch design consisting of three cycles of 1-minute OFF and 1-minute ON periods. Photoactivation of CeA^Isl1^ neurons significantly increased the time and frequency that mice spent biting food (Fig. 3, G to I) or Styrofoam (Fig. 3, L to N), compared to both OFF epochs and control mice. The same effect was observed in mice after 4 hours of food deprivation, suggesting that the biting behavior was not much influenced by internal state (fig. S4, A to C). In addition to object-directed biting, photoactivation of CeA^Isl1^ neurons also induced robust fictive feeding behavior in the absence of edible (food) (Fig. 3, J and K, fig. S4, D and E) or inedible (Styrofoam) stimuli (Fig. 3, O and P). Fictive feeding behaviors were characterized by forepaw-to-mouth movements resembling consumption. These behaviors were spatially biased: fictive feeding was significantly more likely to occur at greater distances from the object compared to object-directed biting (Fig. 3, Q and R). These observations suggest that activation of CeA^Isl1^ neurons is sufficient to engage the core motor programs underlying feeding behavior, even in the absence of external sensory cues or ingestible material.

To test whether CeA^Isl1^ neurons influence food consumption, we measured food intake in fed mice using both optogenetic and chemogenetic activation. For optogenetics, mice received blue light stimulation in alternating 20-minute ON/OFF epochs, and food intake was averaged across conditions (fig. S4, F and G). For chemogenetic activation, CNO was administered 30 minutes before testing, and food intake was measured at 0.5, 1, 2, and 3 hours (fig. S4, H and I). In both cases, activation of CeA^Isl1^ neurons did not significantly alter total food intake in sated animals.

Our calcium imaging experiments revealed that CeA^Isl1^ neurons were strongly activated during gnawing on crickets. To test whether this activity pattern reflected a causal role in predatory behavior, we performed a cricket hunting assay in mice food-deprived for 4 hours to increase motivation. Mice were presented with three live crickets, and blue light (473 nm) was delivered in an alternating 1-minute OFF /1-minute ON design for three cycles. Consistent with our imaging data, optogenetic activation did not alter the time spent investigating or pursuing crickets (fig. S4, J and K), but significantly increased the time spent capturing prey and engaging in fictive feeding in the absence of crickets (fig. S4, L and M). Similarly, in a novel object exploration task, CeA^Isl1^ neurons stimulation did not increase the time spent near or frequency of visits to the novel object (fig. S4, P to R), indicating that activation does not enhance motivation for exploration or novelty-seeking behavior. Notably, the time spent eating and the total number of crickets killed did not differ between ChR2 and control (EGFP) mice (fig. S4, N and O), suggesting that photoactivation of CeA^Isl1^ neurons promotes bite-related behaviors without enhancing overall predatory efficiency.

### CeA^sl1^ neurons are necessary for efficient biting of solid food

Having established that activation of CeA^Isl1^ neurons enhanced vigor of orofacial movements during linguine consumption, we next asked whether these neurons were necessary for executing efficient biting. To test this, we optogenetically inhibited CeA^Isl1^ neurons using archaerhodopsin (Arch) while mice engaged in a solid food consumption assay (Fig. 3S). We first challenged the mice with uncooked linguine. During light-induced inhibition (5321⍰m), mice showed significantly prolonged feeding bouts compared to Arch-OFF and EGFP controls (Fig. 3, T and U), while the frequency of feeding bouts remained unchanged (fig. S5A). This suggested that CeA^Isl1^ neuronal activity was not required to initiate feeding per se, but was critical for sustaining efficient oromotor output. Notably, total consumption remained unchanged (fig. S5, B and C), indicating a disruption in execution rather than appetite. To determine whether this disruption depended on food hardness, we next used a more fragile substrate of similar taste—uncooked spaghetti with a smaller diameter (1 mm) (fig. S5D). In this condition, CeA^Isl1^ neuron inhibition in fasted mice led to a partial disruption in feeding efficiency, with significantly longer consumption times observed for the second pasta piece in Arch-ON mice (fig. S5, E to I). These results suggested that CeA^Isl1^ neurons were necessary for efficient biting behavior which became most obvious when mice processed resistant food items or when they reached satiety. When mice were tested with softer food, such as standard chow pellets, no impairment was observed (fig. S5, J to O).

To examine how CeA^Isl1^ neuron activity influences orofacial motor output, we recorded electromyographic (EMG) activity from two jaw closing muscles, masseter and temporalis muscles using flexible Myomatrix arrays^32^ implanted in awake, freely moving mice (Fig. 3V). In Isl1-CreER mice expressing the inhibitory DREADD hM4Di in CeA^Isl1^ neurons, food-biting behavior generated robust EMG signals under saline conditions (Fig. 3W). Chemogenetic inhibition of CeA^Isl1^ neurons with CNO markedly reduced the amplitude of EMG bursts in both muscles (Fig. 3, W to Y), indicating that CeA^Isl1^ neurons activity is required for normal recruitment of jaw-closing motor units during feeding.

We next tested whether the contribution of CeA^Isl1^ neuron activity to jaw-muscle recruitment depended on the mechanical properties of the object being bitten. Chemogenetic inhibition of CeA^Isl1^ neurons markedly reduced masseter and temporalis EMG amplitudes during biting of harder substrates, including softwood and Styrofoam (fig. S5, P to S). By contrast, the effect was attenuated for softer materials: soft silicone produced only partial reductions (fig. S5, T and U), and sponge biting showed minimal or no change (fig. S5, V and W). These results indicate that CeA^Isl1^ neuron activity is preferentially required for generating high-force jaw closure, with weaker engagement during low-effort biting of compliant materials.

### Activation of CeA^Isl1^neurons supports reinforcement and reward-associated learning

Neurons in the CeA have previously been shown to be activated by both aversive and appetitive stimuli^11,33^, raising the question of whether CeA^Isl1^ neurons contribute to the positive or negative motivational value of experience. We used real-time place preference (RTPP) to test whether CeA^Isl1^ neurons supported positive reinforcement. During testing, ChR2-expressing mice showed a strong preference for the light-paired compartment, which reversed when stimulation contingencies were switched. Control mice showed no such preference, indicating that CeA^Isl1^ neuron activation is intrinsically reinforcing (Fig. 4, A and B). To further test the reinforcing properties of CeA^Isl1^ neurons, we conducted an intracranial self-stimulation (ICSS) assay, in which mice nose-poked for bilateral CeA^Isl1^ neuron photostimulation. ChR2 mice rapidly developed robust ICSS behavior, whereas EGFP controls did not (Fig. 4C), confirming that CeA^Isl1^ neuronal activation is sufficiently rewarding to support operant reinforcement.

**Figure 4.**
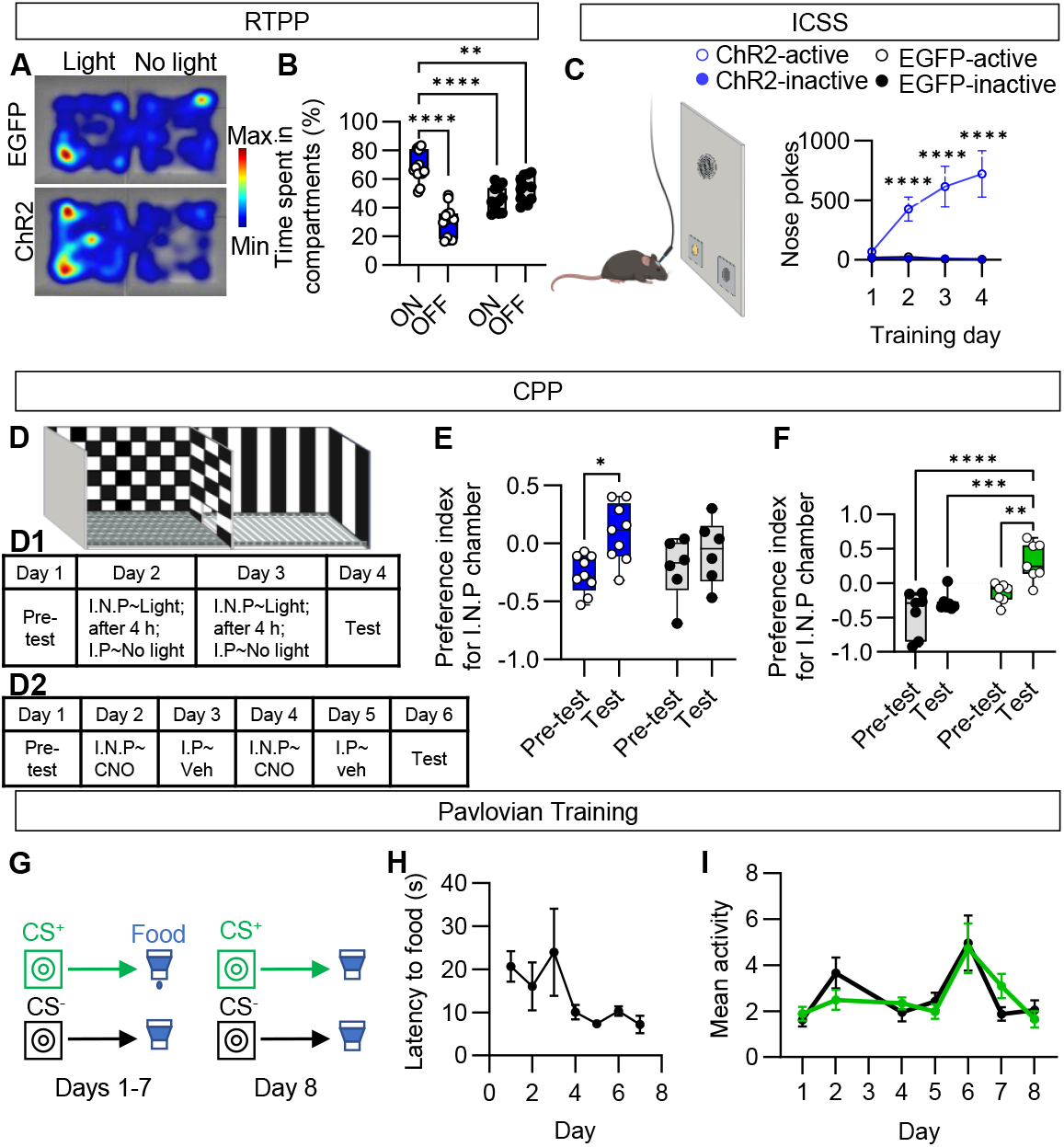
Activation of CeA^Isl1^ neurons supports positive reinforcement and conditioned reward learning. **A**. Heatmaps showing representative occupancy plots of EYFP (top) and ChR2 (bottom) mice. **B**. Quantification of time spent in the light-paired (ON) and non-paired (OFF) compartments (n = 13 mice per group; ****p < 0.0001, two-way ANOVA with Sidak’s multiple comparisons). **C**. Intracranial self-stimulation assay (ICSS). Nose poke behavior across four days of training in ChR2 (blue) and EGFP (black) mice (n = 11 mice per group; ****p < 0.0001, two-way ANOVA with Sidak’s multiple comparisons). **C**. Schematic of CPP chamber. Initially preferred (I.P.) chamber and initially non-preferred (I.N.P.) chamber. **D1**. Optogenetic assay: mice underwent a pre-test (Day 1), two days of conditioning (light-paired I.N.P. or unpaired I.P. compartments); and a test session without stimulation (Day 4). **D2**. Chemogenetic activation assay: mice underwent a pre-test (Day 1). On alternating days (Day 2 and 4), mice received CNO paired I.N.P. compartment), while on Day3 and 5, they received vehicle paired with I.P. compartment, and a test session (Day 4). **E**. Preference index toward I.N.P. chamber on test versus pre-test days in ChR2 (blue) and control (grey) mice. N = 9 mice per group; *p < 0.05; two-way ANOVA with Sidak’s multiple comparisons. **F**. Preference index toward I.N.P. chamber on test versus pre-test days in hM3Dq (green) and control (grey) mice. (n = 7 mice per group; **p < 0.01, ***p < 0.001, ****p < 0.0001; two-way ANOVA with Sidak’s multiple comparisons). **G**. Schematic of Pavlovian training protocol. From Days 1–7, a conditioned stimulus (CS^+^) was paired with food delivery, while a second stimulus (CS^−^) was never paired. Day 8 served as a probe test. **H**. Latency to retrieve food (n = 7 mice). **I**. Mean CeA^Isl1^ neuron activity in response to CS^+^ (green) and CS^−^ (black) cues across training days.

Natural rewards not only reinforce instrumental behavior but also support Pavlovian learning, where predictive cues acquire incentive value and promote approach even in the absence of the reward ^34^. We next asked whether CeA^Isl1^ neuron activation could drive conditioned place preference (CPP). In this assay, ChR2-expressing mice developed a significant preference for the light-paired context, and similarly, chemogenetic activation of CeA^Isl1^ neurons led to a significant preference for the CNO-paired context, while control mice did not show a preference in either condition (Fig. 4, D and F), suggesting that CeA^Isl1^ neuron activation is sufficient for forming reward–context associations. Finally, to assess whether CeA^Isl1^ neurons encoded the predictive value of cues during reward learning, we performed in vivo calcium imaging during a Pavlovian conditioning task. Mice were trained to associate one cue (CS^+^) with food delivery and another (CS^−^) with no outcome (Fig. 4G). Behaviorally, animals learned the association, showing a progressive reduction in latency to retrieve the reward across training days (Fig. 4H). However, CeA^Isl1^ neuronal activity did not differ between CS^+^ and CS^−^ trials, even after learning (Fig. 4I), suggesting that while these neurons can drive reinforcement and support associative learning, they do not encode cue-specific predictive value during Pavlovian conditioning.

### Distinct CeA–brainstem pathways drive biting behavior

To begin addressing the circuit mechanism by which CeA^Isl1^ neurons control biting behavior, we anatomically mapped the long-range outputs of CeA^Isl1^ neurons. We found that they projected to hindbrain targets including the PCRt, PBN, PPTg, nucleus of the solitary tract (NTS), microcellular tegmental nucleus (MiTg), and supratrigeminal nucleus (Su5) (Fig. 5, A and B). These target regions have all be shown to be directly or indirectly involved in motor control. The PCRt is involved in coordination of orofacial movements and autonomic functions^14,35^. The PBN is a critical meal-termination center that integrates visceral and nociceptive signals to suppress feeding and promote aversion^36,37^. The PPTg regulates locomotion, playing a key role in movement initiation and maintenance^38,39^. The NTS receives visceral afferents from the vagus nerve and drives reflexive or patterned motor responses, such as swallowing, gagging, or vomiting^39,40^. The MiTg is likely a premotor brainstem nucleus involved in coordinating specific motor patterns^41^ and the Su5 provides direct premotor control of orofacial motor neurons^42,43^.

**Figure 5.**
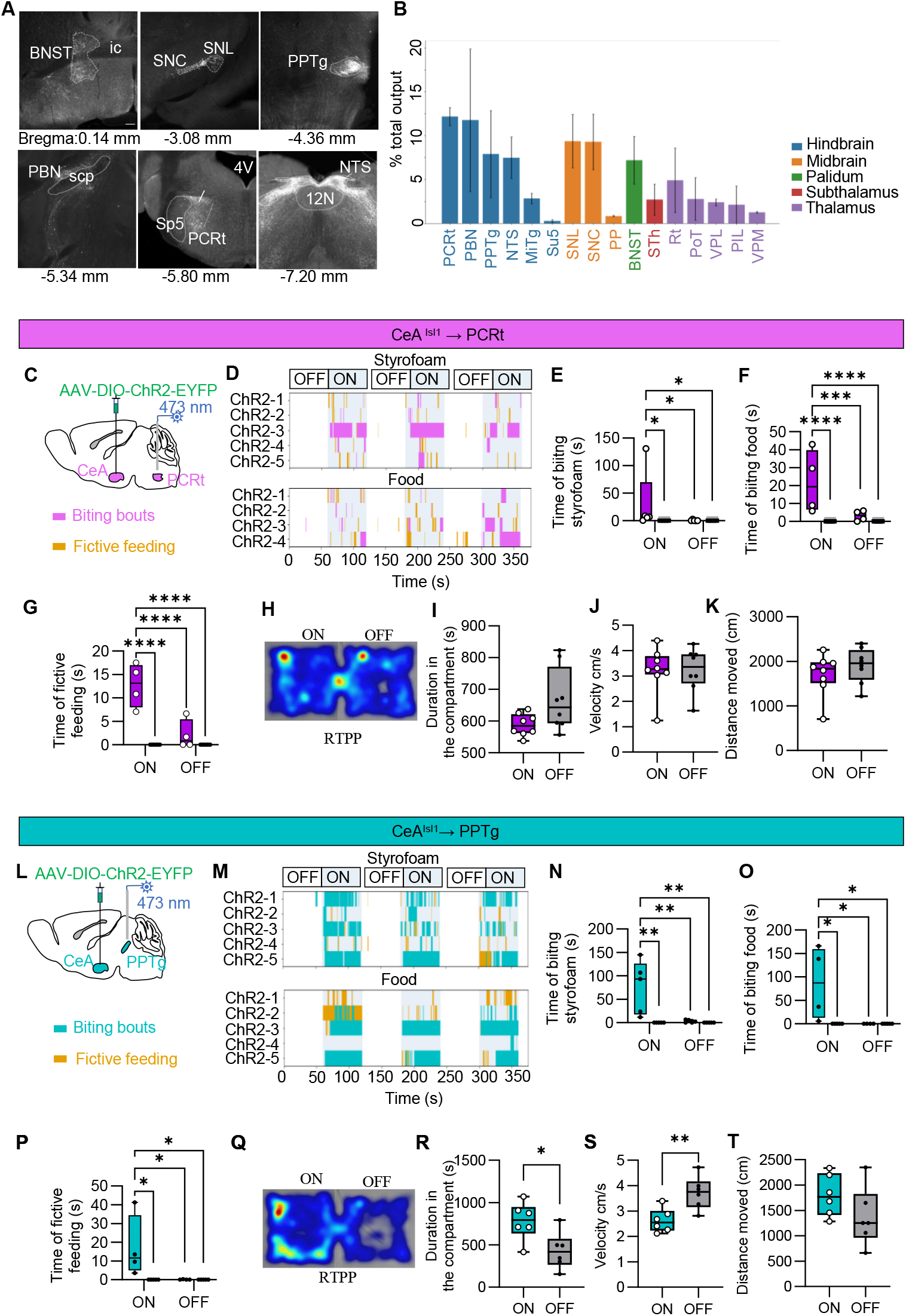
CeA^Isl1^ → PCRt terminal stimulation promotes object-directed biting behavior. **A**. Representative images of brain areas innervated by CeA^Isl1^ neurons. Scale bar, 100 µm. **B**. Major brain regions innervated by CeA^Isl1^ neurons depicted as percentage (mean ± s.e.m.) of total outputs (n = 3 mice). **C**. Schematic showing viral targeting of CeA^Isl1^ neurons and terminal stimulation in the PCRt using 473 nm light in Isl1-CreER mice. **D**. Raster plot showing biting bouts (pink) and fictive feeding bouts (orange) across individual ChR2 mice during alternating light OFF and light ON epochs in biting Styrofoam and food assay. **E-G**. Quantification of behaviors during ON and OFF periods for both ChR2 (pink) mice and EGFP (grey) mice in biting assay. **E**. Duration of biting Styrofoam. **F**. Duration of biting food. **G**. Duration of fictive feeding. N = 5 mice per group. Two-way ANOVA with Tukey’s post hoc test, *P1<10.05, ***P1<10.001, ****P1<10.0001. **H**. Representative heatmap showing occupancy plots of CeA ^Isl1^ → PCRt ChR2 mice. **I**. Quantification of time spent in the light-paired (ON) and non-paired (OFF) compartments (n = 5 mice per group; N.S., two-tailed unpaired t-test). **J**. Average velocity in the light-paired (ON) and non-paired (OFF) compartments (n = 5 mice per group; N.S., two-tailed unpaired t-test). **K**. Total distance moved in the light-paired (ON) and non-paired (OFF) compartments (n = 5 mice per group; N.S., two-tailed unpaired t-test). **L**. Schematic showing viral targeting of CeA^Isl1^ neurons and terminal stimulation in the PPTg using 473 nm light in Isl1-CreER mice. **M**. Raster plot showing biting bouts (green) and fictive feeding bouts (orange) across individual ChR2 mice during alternating light OFF and light ON epochs in biting Styrofoam and food assay. **N-P**. Quantification of behaviors during ON and OFF periods for both ChR2 (green) mice and EGFP (grey) mice in biting assay. **N**. Duration of biting Styrofoam. **O**. Duration of biting food. **P**. Duration of fictive feeding. N = 5 mice per group. Two-way ANOVA with Tukey’s post hoc test, *P ⍰< ⍰ 0.05, **P ⍰ < ⍰ 0.01. **Q**. Representative heatmap showing occupancy plots of CeA^Isl1^ → PPtg ChR2 mice. **R**. Quantification of time spent in the light-paired (ON) and non-paired (OFF) compartments (n = 5 mice per group; *p < 0.05, two-tailed unpaired t-test). **S**. Average velocity in the light-paired (ON) and non-paired (OFF) compartments (n = 5 mice per group; **p < 0.01, two-tailed unpaired t-test). **T**. Total distance moved in the light-paired (ON) and non-paired (OFF) compartments (n = 5 mice per group; N.S., two-tailed unpaired t-test). Abbreviations: BNST, the bed nucleus of the stria terminalis; MiTg, microcellular tegmental nucleus; NTS, nucleus of the solitary tract; PAG, periaqueductal gray; PBN, parabrachial nucleus; PCRt; parvicellular reticular nucleus; PP, peripeduncular nucleus; PPTg, pedunculopontine tegmental nucleus; PIL, posterior intralaminar thalamic nucleus; PoT, posterior thalamic nuclear group, triangular Part; Rt, reticular thalamic nucleus; SNL, substantia nigra, lateral part; SNC, substantia nigra, compact part; STh, subthalamic nucleus; Su5, supratrigeminal nucleus; scp superior cerebellar peduncle (brachium conjunctivum); VPL, ventral posterolateral thalamic nucleus; VPM, ventral posteromedial thalamic nucleus.

In addition, we found that CeA^Isl1^ neurons projected to other forebrain and midbrain targets including the subthalamic nucleus (STh), substantia nigra pars lateralis (SNL) and pars compacta (SNC), the peripeduncular nucleus (PP), the bed nucleus of the stria terminalis (BNST), the reticular nucleus of the thalamus (Rt), the posterior triangular thalamic nucleus (PoT), the ventral posterolateral (VPL), posterior intralaminar (PIL), and ventral posteromedial (VPM), thalamic nuclei (Fig. 5, A and B). These regions are known to be involved in motor regulation (STh, SNL, SNC)^44,45^, reward and arousal (BNST, SNC)^46^, sensorimotor integration (PP, PoT, VPL, VPM, PIL)^47^, and sensory gating or attentional filtering (Rt)^48,49^. Together, these projection patterns are consistent with a model that CeA^Isl1^ neurons coordinate orofacial motor programs and visceral reflexes with broader behavioral state and sensorimotor context.

To dissect the downstream circuits by which CeA^Isl1^ neurons regulate biting behavior, we examined two major brainstem projections: the CeA^Isl1^ → PCRt and the CeA^Isl1^ → PPTg pathways. To investigate the CeA^Isl1^ → PCRt projection, we injected AAV-DIO-ChR2-EYFP into the CeA of Isl1-CreER mice and implanted optic fibers above the PCRt (Fig. 5C). Mice were presented with five non-edible Styrofoam objects that elevated calcium signals in the majority of CeA^Isl1^ neurons (Fig. 1). Optogenetic stimulation of CeA^Isl1^ axon terminals in the PCRt significantly increased both the duration and frequency of biting behavior directed toward the Styrofoam (Fig. 5, D and E, fig. S6A). Similar increases in duration and frequency of biting behavior were observed towards pieces of normal chow (Fig. 5, D and F, fig. S6B). We also found robust fictive feeding behavior induced by photoactivation of CeA^Isl1^ → PCRt projection in both cases (Fig. 5G). These results demonstrate that CeA^Isl1^→ PCRt projections are sufficient to drive object-directed orofacial actions, suggesting a functional role for this pathway in promoting biting-related motor programs.

To assess whether CeA^Isl1^ → PCRt stimulation carries motivational valence, we performed RTPP testing (Fig. 5, H to K). Mice did not develop preference for the photostimulation-paired chamber (Fig. 5, H and I), indicating that CeA^Isl1^ → PCRt activation lacks positive reinforcement. Locomotor activity analysis revealed no significant changes in velocity or total distance traveled (Fig. 5, J and K), demonstrating that the behavioral effects were not due to general arousal or hyperactivity. Together, these data identify the CeA^Isl1^ → PCRt pathway as a motor-promoting circuit that executes the act of biting without encoding motivational value.

Stimulation of CeA^Isl1^→ PPTg projections (Fig. 5L) also produced robust increases in biting toward both inedible and edible objects, as well as fictive feeding, similar to the CeA^Isl1^→ PCRt pathway (Fig. 5, M to P, fig. S6, C and D). However, in the RTPP assay, the CeA^Isl1^→ PPTg pathway was positively reinforcing, as mice spent significantly more time in the light-paired compartment (Fig. 5, Q and R). These mice also exhibited decreased velocity but unchanged total movement distance (Fig. 5, S and T), suggesting enhanced approach motivation. Together, these findings reveal that CeA^Isl1^ → PCRt and CeA^Isl1^→ PPTg projections constitute distinct components of an emotional–motor network: the CeA^Isl1^ → PCRt projection provides a descending command for bite execution, while the CeA^Isl1^→ PPTg projection energizes and reinforces biting behavior through positive motivational drive.

Next, we aimed at identifying the monosynaptic inputs to CeA^Isl1^ neurons with a focus on the CeA^Isl1^ subpopulation that projects to the PCRt. A Flp-dependent rabies glycoprotein (RVG) and TVA receptor were co-injected into the CeA, while Cre-dependent Flp recombinase was injected into the PCRt and delivered retrogradely via rAAV2-retro into the CeA and other incoming brain areas (fig. S6E). This strategy ensured that only CeA^Isl1^ neurons projecting to the PCRt would express the necessary components for rabies virus infection and trans-synaptic spread. Following rabies virus injection into the CeA, we found the start cells mainly located in the anterior part of CeM (fig. S6F). Input neurons were identified throughout the brain. Quantitative analysis revealed substantial input from several forebrain and brainstem regions. Notably, the strongest sources of input originated from the lateral part of the central amygdala, the basolateral amygdala (BLA), lateral hypothalamus (LH), insular cortex, and several regions of the motor and sensory cortex (fig. S6G). These results suggest that CeA^Isl1^→ PCRt neurons integrate diverse motivational, sensory, and motor-related signals to regulate brainstem motor output.

Building on these circuit- and behavior-level findings, we next examined whether CeA^Isl1^ neurons were conserved across species. Transcriptional analyses revealed that *Isl1*^+^ neurons were present in the CeA or CeA-like regions of human^50,51^, mouse^52^, chicken^51^, lizard^53^, and even salamander^54^ (fig. S7, A to E). We compared *Isl1* with *Prkcd*, a canonical marker of CeA neurons that mediate aversive and defensive behaviors, because this contrast highlights functional diversity within the CeA. In both mouse and human, *Pkc*δ^+^ and *Isl1*^+^ neurons were largely non-overlapping (fig. S7, C to E, H to J), indicating that they represent distinct subtypes. In lizard, only minimal overlap was observed (fig. S7, A and F), whereas in chicken approximately 50% of *PRKCD*^+^ cells also expressed *ISL1* (fig. S7, B and G), suggesting a less specialized CeA organization. In salamander, *Prkcd* expression was absent, and only *Isl1*^+^ cells were detected in the CeA-like region (fig. S7, K to M). Thus, *Isl1* emerges as a highly conserved molecular signature of CeA-like neurons across vertebrates (fig. S7N), underscoring its fundamental role in the evolutionary blueprint of this structure.

## Discussion

Our results uncover a previously unrecognized function of CeA neurons in encoding the physical properties of a bitten object, with neuronal activity scaling proportionally to the object’s hardness. Specifically, we show that stimulation of CeA^Isl1^ neurons not only triggers biting behavior but also disrupts the fine control of bite force, whereas inhibition of these neurons impairs bite efficiency. These findings indicate that CeA^Isl1^ neurons help calibrate jaw motor output to match the physical demands of different objects, ensuring that forceful, adaptive biting is appropriately scaled to substrate hardness.

Our calcium imaging data revealed that edible and non-edible food items with different physical properties (density, viscosity) activated the majority of CeA^Isl1^ neurons to varying degrees. Depending on the item, CeA^Isl1^ neurons were recruited into different ensembles and the harder the object, the stronger the activation, suggesting that these neurons encode the range of physical properties of food items. In contrast, the sensory attributes (taste, smell) of food had minor effects on CeA^Isl1^ neurons, with most flavors eliciting similar activation patterns. These characteristics differentiate CeA^Isl1^ neurons well from other CeA appetitive neurons. CeM^Htr2a^ neurons were previously shown to drive the consumption of food and water and CeM^Sst^ neurons to drive water consumption only^12^. Both populations increase their activities during consumption of rewarding substances. When switching between rewards of different physical attributes (e.g. solid food and liquid Fresubin), the activity of CeM^Htr2a^ neurons often remained stable, suggesting that these neurons pay little regard to the physical attributes of food. Mechanistically, these two populations prolong food or water consumption by conveying stimulus-specific signals to downstream reward centers, rather than promoting biting or licking directly^12,33^.

Calcium imaging further revealed that CeA^Isl1^ neurons are preferentially engaged during gnawing but show minimal responses during subsequent chewing, highlighting a functional distinction in how the CeA modulates orofacial behaviors. This specificity suggests that CeA circuits facilitate motor programs that demand forceful, precise interactions with objects, such as biting and gnawing, rather than the automatic, rhythmic mastication governed by local brainstem central pattern generators. Indeed, previous work has shown that distinct premotor circuits within the supratrigeminal nucleus and trigeminal motor system differentially coordinate gnawing and chewing movements, with gnawing requiring higher jaw-closing forces and incisors, whereas chewing primarily involving rhythmic molar action^55 42,56^. This is in line with older work demonstrating that as the hardness of food increases, both the number of chewing cycles and the level of jaw-closing muscle EMG activity also increase, but that EMG activity progressively decreases during the sequence as the food is broken down^57,58^. This suggests that forceful, adaptive bite control is especially critical during the initial interaction with hard food items, the phase during which CeA^Isl1^ neurons are most active in our study. Once the bolus is broken down, the demands for precise force modulation diminish and chewing proceeds through well-defined rhythmic circuits in the brainstem^56^. Together, these observations support a model in which CeA^Isl1^ neurons couple sensory evaluation of substance properties to the initiation and calibration of bite force, linking motivational state to skilled orofacial motor output.

Our photoactivation experiments showed that CeA^Isl1^ neurons are sufficient to increase bite force and frequency of biting, without increasing food consumption. The activity of CeA^Isl1^ neurons is not much influenced by the hunger state of the animals, in sharp contrast to CeM^Htr2a^ neurons which are activated by the hunger hormone ghrelin^13^. Biting activity towards edible and non-edible items was instead spatially biased: object-directed biting occurred at a distance of approx. 2 cm from the mouth, whereas fictive feeding behavior was favored at larger distances. The circuit properties underlying this spatial bias are currently unclear and will be an interesting future research focus. Photoinhibition experiments showed that the activity of CeA^Isl1^ neurons is necessary for efficient biting of solid food that requires strong bite force, consistent with a role in scaling motor output to match physical demands.

Our circuit analysis revealed that CeA^Isl1^ neurons regulate biting behavior through two parallel output pathways that differentially contribute to motor execution and motivational reinforcement. Activation of the CeA^Isl1^→ PCRt projection robustly drove biting, indicating direct control of orofacial motor circuits. In contrast, stimulation of the CeA^Isl1^→ PPTg pathway not only elicited biting but also produced a reinforcing effect, suggesting recruitment of reward-related mechanisms. The PPTg is a key mesencephalic hub that modulates dopaminergic neurons in the substantia nigra and ventral tegmental area. Previous work has shown that PPTg stimulation evokes striatal dopamine release via cholinergic and glutamatergic transmission^59-61^, supporting the idea that the CeA^Isl1^→ PPTg projection may link emotional or sensorimotor signals from the amygdala to midbrain reward systems. CeA^Isl1^→ PPTg stimulation also reduced locomotor velocity despite its reinforcing properties. This reduction likely reflects a behavioral shift from exploration to focused consummatory engagement—a hallmark of reward anticipation and goal-directed approach states—rather than motor suppression. Conversely, the CeA^Isl1^→ PCRt pathway likely mediates the direct motor component of biting through reticular premotor circuits controlling jaw and tongue movements. Together, these results uncover a dual output architecture in which the CeA orchestrates both the kinematics and motivation of orofacial actions. This organization provides a neural substrate through which emotional states can dynamically transform simple motor patterns—like biting—into goal-directed and affectively charged behaviors.

Understanding the afferent inputs to distinct CeA cell populations is essential for deciphering how different neuron types integrate sensory, emotional, and motivational information to orchestrate behavior. For example, CeA^Htr2a^ neurons, which are linked to hedonic feeding, receive strong input from homeostatic and reward-related regions such as the arcuate nucleus (Arc), parasubthalamic nucleus (PSTN), substantia nigra pars lateralis (SNL), and dorsal raphe nucleus (DR) ^8^. In contrast, CeA^Isl1^ neurons show minimal input from these classic feeding and neuromodulatory centers but instead receive robust inputs from limbic and associative areas including the basolateral amygdala (BLA), medial amygdala (MeA), ventral pallidum (VP), piriform cortex (Pir), and caudate putamen (CPu). This pattern suggests that CeA^Isl1^ neurons may play a broader role in integrating valence, social, and motivational signals. Notably, the strong input from the MeA, which is absent for CeA^Htr2a^ neurons, positions CeA^Isl1^ neurons to process social or pheromonal cues that could shape behaviors such as differentiating aggressive bites from parental pup retrieval. Similarly, substantial input from the interstitial nucleus of the posterior limb of the anterior commissure (IPAC) implies that CeA^Isl1^ neurons may integrate internal state signals like stress and reward ^62-64^to flexibly adjust behavioral outputs, for example, modulating feeding or hunting in response to threat. Future work combining precise circuit mapping with behavioral testing will be crucial to reveal how these inputs shape specific motivated behaviors.

The extensive projections of CeA^Isl1^ neurons to multiple brainstem and forebrain regions underscore the CeA’s role as a hub that dynamically integrates motivational and sensory signals with premotor circuits. For example, projections to brainstem areas such as the PBN, NTS, and Su5 likely facilitate the coordination of orofacial motor programs with visceral feedback^36,42,65^. Inputs to the PPTg and substantia nigra may further modulate arousal, locomotor drive, and reinforcement learning, thereby linking emotional or motivational states to action selection and behavioral vigor^66,67^. Projections to the BNST and PP could contribute to sustained affective, social, and defensive states by integrating multimodal sensory and motivational information to modulate anxiety-like behaviors, maternal responses, social interaction, and context-dependent threat or arousal reactions ^68,69^. Meanwhile, connections to multiple thalamic relay nuclei, including somatosensory (VPL, VPM) and polymodal (PIL, PoT) regions, raise the possibility that the CeA exerts top-down modulation over sensory gain, biasing perception in a state-dependent manner to promote or suppress specific behavioral repertoires^70,71^. Future work will be needed to dissect how specific CeA^Isl1^ subpopulations differentially engage these downstream targets to coordinate discrete aspects of orofacial and locomotor behaviors. Moreover, functional experiments targeting the interactions between CeA outputs to motor nuclei and ascending sensory relays could elucidate how motivational circuits override or reshape sensory processing to support goal-directed actions. This distributed output architecture may be especially relevant for understanding how emotional states contribute to maladaptive conditions such as compulsive gnawing, pica, or stress-induced sleep bruxism.

## Acknowledgements

We thank Sofia delgado for help with management of the animal colony; Helena Weltzien for help with behavior experiment; Yue Zhang (Department of Synapses – Circuits – Plasticity, Max-Planck Institute for Biological Intelligence) for help with data analysis; Karl-Klaus Conzelmann (Gene Center Munich, LMU) for providing EnvA G-deleted rabies virus. The Myomatrix Array usage reported in the publication was supported by the NIH BRAIN Initiative under award numbers NIH U24NS126936 and NIH R01NS109237. A.P.A. was supported by FAPESP fellowship no. 2023/02896-3. This study was supported by the Max-Planck Society and the European Research Council under the European Union’s Horizon 2020 research and innovation programme (no. 885192, BrainRedesign).

## Author contributions

WD and RK conceptualized and designed the study. WD conducted the experiments and analyzed data. AP assisted with surgery and behavior experiments. WK and CQ assisted with data analysis. WD and RK wrote the paper with input from all authors. R.K. supervised and provided funding.

## Data Availability

The data supporting the current study are available from the corresponding author upon reasonable request.

## Materials and methods

### Animals

Male and female mice that were at least 2 months old were used in all experiments, following regulations from the government of Upper Bavaria. The mice were housed in their home cages with 2-5 mice per cage, under a 12-hour light/12-hour dark cycle, with food and water freely available. The Isl1-CreER transgenic line (Isl1^tm1 (cre/Esr1*) Krc^/SevJ) was purchased from Jackson Laboratories. Prkcd-Cre (Tg (Prkcd-glc-1/CFP, -Cre) EH124Gsat) BAC mice were imported from the Mutant Mouse Regional Resource Center (https://www.mmrrc.org/). Ai9 (B6. Cg-Gt (ROSA) 26Sor^tm9 (CAG-tdTomato) Hze^/J) mouse line was as described previously, Transgenic mice were bred onto a C57BL/6NRj background (Janvier Labs - http://www.janvier-labs.com). For optogenetic and chemogenetic manipulations, animals were handled and housed singly on a 12-hour inverted light cycle for at least 3 days prior to the experiments. Except during food deprivation for feeding experiments, mice were given ad libitum food access. All behavior tests were carried out at the same time each day during the dark period (1 p.m.–6 p.m.).

### Viral vectors

The following AAV viruses were obtained from Addgene: AAV9-pAAV-hSyn-DIO-hM3D (Gq)-mCherry (Addgene, 44361), pAAV-hSyn-DIO-hM4D (Gi)-mCherry (Addgene, 44362), AAV2-pAAV-hSyn-DIO-mCherry (Addgene, 50459), AAV5-Ef1a-DIO-ChR2-EYFP (Addgene, 35509), AAV5-Ef1a-DIO-EGFP (Addgene, 27056). AAV5-EF1a-DIO-eArch3.0-EYFP were produced at the Gene Therapy Center Vector Core at the University of North Carolina Chapel Hill. pssAAV-2-hSyn1-dlox-jGCaMP8m(rev)-dlox-WPRE-SV40p(A) (v628-1) was purchased from Viral Vector Facility (VVF) in Neuroscience Center Zurich (ZNZ) at the University of Zurich and ETH Zurich. rAAV2/8-nEf1α-FDIO-RVG-WPRE-hGH polyA, rAAV2/8-nEF1α-fDIO-EGFP-T2A-TVA-WPRE-hGH polyA, and rAAV2 (Retro)-EF1α-DIO-Flp-WPRE-hGH polyA were produced by BrainVTA (Wuhan). EnvA G-deleted rabies for long-range monosynaptic tracing was a gift from Karl-Klaus Conzelmann (Gene Center Munich, LMU).

### Stereotactic surgery

Mice were anesthetized with 1.5-2% isoflurane and received oxygen at 1.0 liter per minute before being placed in a stereotaxic frame (Kopf Instruments). A heating pad was used to keep the body temperature stable. Carprofen (5 mg/kg bodyweight) was used subcutaneously as an analgesic.

Once the mouse skull was exposed, we drilled a cranial window (1–2 mm^2^) unilaterally (calcium imaging experiments, monosynaptic rabies and anterograde tracing experiments) or bilaterally (optogenetic and chemogenetic experiments). Next, a glass pipette (#708707, BLAUBRAND intraMARK) was lowered into the window to deliver 300 nl of viral vector to the area of interest (coordinates: CeA: −1.22 mm anterior to bregma, ± 2.6 mm lateral from midline and, −4.85 mm vertical from the brain surface; PCRt: − 5.6 mm anterior to bregma, ± 1.27 mm lateral from midline and, − 4.7 mm vertical from the brain surface; PPTg: − 4.7 mm anterior to bregma, ± 1.2 mm lateral from midline and, − 3.5 mm vertical from the brain surface). In the same surgery, mice used in optogenetic experiments were bilaterally implanted with optic fibers (200 µm core, 0.39 NA, 1.25-mm ferrule (Thorlabs)) above the CeA (− 4.35 mm ventral), the PCRt (− 4.2 mm ventral), or the PPTg (-3.0 mm ventral). Implants were secured with cyanoacrylic glue, and the exposed skull was covered with dental acrylic (Paladur). For all other mice, the incision was closed with sutures.

Isl1-CreER mice used for rabies tracing were first unilaterally injected in the PCRt with rAAV2 (Retro)-EF1α-DIO-Flp and in the CeA with the starter rAAV2/8-nEf1α-FDIO-RVG and rAAV2/8-nEF1α-fDIO-EGFP-T2A-TVA. After 4 weeks, the same mice were injected with the rabies virus. Seven days later, mice were killed, and their brain tissue was collected and processed for immunohistochemistry.

### GRIN lens implantation and baseplate fixation

Four weeks after GCaMP8m viral injection in the CeA, mice were implanted with a gradient index (GRIN) lens. At the same coordinates of the injection, a small craniotomy was made and a 23G needle was slowly lowered into the brain to clear the path for the lens to a depth of -4.5 mm from bregma. After retraction of the needle, a GRIN lens (ProView lens; diameter, 0.6 mm; length, ∼7.3 mm, Inscopix) was slowly implanted above the CeA and then fixed to the skull using UV light-curable glue (GRADIATM DIRECT FLO) and iBOND® Universal (Kulzer). 4 weeks after GRIN lens implantation, mice were “baseplated” under anesthesia. Briefly, in the stereotaxic setup, a baseplate (BPL-2; Inscopix) was positioned above the GRIN lens, adjusting the distance and the focal plane until the neurons were visible. The baseplate was fixed using UV light-curable glue (GRADIATM DIRECT FLO) C…B Metabond (Parkell). A baseplate cap (BCP-2, Inscopix) was left in place to protect the lens.

### Tamoxifen protocol

Tamoxifen solution was prepared in 90% corn oil and 10% ethanol. The injection dose was determined by weight (using approximately 200 mg tamoxifen/kg body weight) and was given for 2 consecutive days. For all AAV expression experiments, tamoxifen was administered 1 day after stereotactical AAV injection.

### Pharmacological treatments

For chemogenetic experiments, mice were given an intraperitoneal (IP) injection of CNO (2 mg/kg for hM3D (Gq) and 0.4 mg/kg for hM4D (Gi) diluted in saline) or the equivalent volume of saline before the experiment and were allowed to recover in their home cages for 30 minutes.

### Optogenetic manipulations

Mice were bilaterally tethered to optic fiber patch cords (Prizmatix) connected to a multi wavelength LED (Prizmatix) via a rotary joint (Prizmatix) and mating sleeve (Thorlabs). For photoactivation experiments, 5 ms, 473-nm light pulses at 20 Hz and 10-15 mW were used. Constant 532-nm light at 18 mW was used for photoinhibition experiments. The LED were triggered, and pulses were controlled with EthoVision XT16 software (Noldus Information Technologies).

### Feeding behavior

Mice were habituated to the behavioral context for daily 10-min sessions, for 2 days before the experiment. For 4h-fasted feeding test, mice were food restricted for 4 hours. Mice were presented with a regular food pellet, and allowed to feed. The weight of the food pellet, including the food debris left in the cage floor after test, was measured to calculate the food intake. For ’fed’ feeding test, mice were not food deprived before the test. In the optogenetic experiment, the light was started just after the mice were put into the testing cage for 20 minutes, then the light was off for 20 minutes. The food intake was measured for both periods. For the pharmacological experiment, CNO were injected 20-30 minutes before the test. The food intake was measured for 0.5 h, 1 h, 2 h and 3 h. All of the feeding tests were performed between 1 pm to 6 pm.

### Optical intracranial self-stimulation

Behavioural training and testing occurred in mouse operant chambers (21.6 cm W x 18.6 cm D x 12.7 cm H, Med Associates Inc.) interfaced with optogenetic stimulation equipment. Prior to the first session, mice were familiarized with purified food pellets (20 mg, TestDiet), which is slightly sweet, in their home cage for 2 consecutive days. Mices were food restriction by time-limiting the availability of food (1 to 2 h) in the home cages (85 % to 90 % of their free-feeding bodyweight) to promote apparatus exploration. Mice explored operant boxes containing an active and inactive nosepoke. On the pre-training day, both active and inactive ports were baited with food treats to encourage exploration. For this study, appetitive stimuli would have impacted latency measurements on the pre-training day and were not used. Training Session length was 60 min, during which time mice were free to respond at any nosepoke port, no food treats were delivered. The inactive nosepoke produced no result. The active nosepoke delivered a 20-Hz train of 5 ms pulses of bluelight for 3 s, the active nosepoke was accompanied by a cue light above the nosepoke and a tone (3k Hz, 5 s), which were insufficient to promote nosepoking in the absence of optical stimulation. Testing occurred once per day for 4 days and port/frequency assignment was counterbalanced. Nosepoke activity was recorded with Noldus EthoVision XT software and visually monitored via infrared camera.

### Real-time place preference test

Mice with optic fiber patch cables tethered were allowed to explore a two-compartment arena (50 cm × 25 cm × 25 cm). Mice were tested across two sessions. In session one, one side was assigned as the photostimulation chamber. Every time the mouse entered this chamber, 5-ms, 473-nm light pulses at 20 Hz and 10-15 mW (measured at the tip of optic fibers) were delivered intracranially for activation experiment. Photostimulation ceased when the mouse left the photostimulation side. In the second session, we assigned the other chamber as the photostimulation side and repeated testing. The behavior of the mice was recorded using a camera. Ethovision XT16 software (Noldus Information Technologies) was used to deliver light pulses and analyze behavioral parameters.

### Conditioned place preference test

A two-compartment CPP apparatus (35 cm L x 18cm W x 29 cm H, connected by a closable door 5 cm x 5 cm) was used for the CPP test. The two compartments have distinct wall patterns and floor grids. Animal location was tracked, and the time spent in each box was assessed with Ethovision XT 16 software (Noldus). For optogenetic experiments: The CPP test consisted of 4 days. On Day 1, each mouse was allowed to explore the entire apparatus freely for 15 min (Pre_Test). After the pre-test the initial preference of each mouse for a given side compartment was recorded. With our apparatus design, most of the mice showed an initial preference for one of the compartments. On days 2 and 3, we placed the mice into their initially non-preferred compartment and delivered 20-Hz light stimulation for 20 min. Approximately 4 h later, we placed the mice into the other side of the compartment without light stimulation. On day 4, we placed the mice back into the apparatus with both compartments accessible for 15 min (without light stimulation) and the time the mouse spent in the compartment is calculated.

For chemogenetic experiments: The CPP test consisted of 6 days. On Day 1, each mouse was allowed to explore the entire apparatus freely for 15 min (Pre_Test). After the pre-test the initial preference of each mouse for a given side compartment was recorded. With our apparatus design, most of the mice showed an initial preference for one of the compartments. Conditioning was initiated on day 2 and encompassed four sessions performed on four consecutive days. In the first conditioning session, mice were injected i.p. with CNO (2 mg/kg) and placed for 1 h in the initially non-preferred (I.N.P.) compartment. On day 3, during the second conditioning session, mice were injected with vehicle (2 % DMSO) and confined for 1 h in the opposite (that is, initially preferred, I.P.) compartment. On day 4 and day 5 the first and second conditioning sessions were repeated, respectively. On day 6, the mice were tested for their side compartment preference by placing them in the left compartment and allowing them to explore the entire apparatus freely for 30 min (post-test). Animal location was tracked, and the time spent in each box was assessed with Ethovision XT 16 software (Noldus). Preference indices (P.I.) were calculated as [(Time in Initially Preferred (I.N.P.) chamber − Time in Initially Preferred (I.P.) chamber)/Total time].

### Pavlovian conditioning

Training was conducted in mouse operant chambers (21.6 cm W x 18.6 cm D x 12.7 cm H, Med Associates Inc.). The chamber had a pellet dispenser that delivered one 20-mg pellet into the port when triggered. This pellet dispenser can monitor the head access by infrared detectors. The chamber contained a multipurpose sound generator (ENV-230, Med Associates Inc), which delivered either a single clear tone (”tone“) or click noise (“click”) when activated. Both stimuli were presented at 75 dB. All conditioned stimuli were 5 s in duration, separated by a variable intertrial interval (ITI) with a mean of 2 min (range = 2 to 6 min). The two auditory stimuli described were used in the experiments; whether the ”click“ or ”tone“ stimulus was the CS+ was counterbalanced across mice. Mice were first food-restricted to 90 % of their baseline body weight and habituated to sucrose pellets (TestDiet 5TUT) in their home cage for 3 days. On the first day of behavioral training, mice learned to retrieve pellets from the food dispenser. During this session, mice received fifteen 20-mg sucrose pellets across a 30-min period. After food port training, mice received conditioning sessions, which lasted 7 days before the imaging test session. Each session lasted 30 min and included 10 presentations of each CS presented in a pseudorandomized order and separated by a variable ITI. During these sessions, termination of one of the cues was followed 1 s later by delivery of one sucrose pellet; this cue was designated the CS+. The other stimulus was presented alone without food and was designated the CS-. During the Pavlovian appetitive extinction test, identical procedures were followed except the reward was not delivered.

### Cricket hunting

To habituate mice to cricket hunting, each mouse was individually placed in an empty cage with three live crickets daily for three consecutive days. Mice were allowed to freely interact with and consume the crickets; the session ended once all three crickets were eaten. On the test day, mice were tested under mild food restriction (2.5⍰g standard chow per day) to increase motivation. At the beginning of each trial, three crickets were introduced into the cage near one corner, while the mouse was positioned in the diagonally opposite corner. For optogenetic stimulation experiments, light was delivered in alternating cycles of 1-minute stimulation followed by 1-minute no stimulation, for a total duration of 6 minutes. For calcium imaging experiments, recording continued until the mouse consumed the final cricket, allowing capture of complete neural dynamics throughout the hunting bout.

### Single-Unit EMG recording

For implantation of the EMG electrodes, the fur over the head and one cheek is shaved. The scalp is disinfected, and the skin is retracted to expose the skull. Small incisions are made above the masseter and temporalis muscles. As much fascia as possible is removed from the muscle, and the target site for implantation is identified. Using blunt forceps, a subcutaneous tunnel is created from the face to the head/neck area of the animal. The electrode leads of myomatrix arrays (RF-4×8-BVS-5) are carefully threaded through the tunnel until they emerge above the target muscle. They are then anchored at several locations within the muscle—two leads in the masseter and two in the temporalis. The skin is closed around the array, and the connector (Omnetics connector, 13 mm, 1.36 g) is secured to the skull. During behavior test session, connect the Intan headstage to the Omnetics connector. The other end of the wire from the headstage is plugged into the recording board (Open Ephys).

### In vivo calcium imaging of freely moving mice

All in vivo imaging experiments were conducted on freely moving mice. GCaMP8m fluorescence signals were acquired using a miniature integrated fluorescence microscope system (nVista – Inscopix) secured in the baseplate holder before each imaging session. Mice were habituated to the miniscope procedure for 3 days before behavioral experiments for 30 min per day. Settings were kept constant within subjects and across imaging sessions. Image acquisition and behavior were synchronized using the data acquisition box of the nVista Imaging System (Inscopix) through a TTL box (Noldus) connected to the USB-IO box from the Ethovision system (Noldus).

Behavioral recordings were conducted in a transparent acrylic chamber equipped with a 45⍰°C angled mirror positioned at the bottom. This setup allowed unobstructed, bottom-up video capture of the mouse’s behavior via a camera positioned below the chamber.

### Object biting

Mice were presented with five distinct objects designed to vary in mechanical properties: softwood, Styrofoam, sponge material, hard silicone, soft silicone, and chow pellet. All materials were cut into standardized cubes (0.5 ⍰ cm × 0.5 ⍰ cm × 0.5 ⍰ cm). Each imaging session lasted 5 minutes, during which the mouse could freely explore and interact with the objects while calcium signals were recorded.

### Flavored milk drinking

For liquid consumption trials, mice were water-deprived for 4 hours prior to testing. Animals remained in their home cage and were sequentially given access to different liquids: strawberry-flavored milk, chocolate-flavored milk, vanilla-flavored milk, water, and 100 ⍰ mM quinine solution. Each liquid was presented for 5 minutes, during which calcium activity and behavioral responses were recorded. For imaging data processing and analysis, we used IDEAS (Inscopix) version 25.1.7.

### Flavored food or sponge biting

Flavored food or sponge were prepared by soaking standard food pellets or sponge cubes in 10 % flavoring solutions (strawberry, vanilla, chocolate), water, or 100 ⍰ mM quinine solution. After drying overnight, the flavored items were presented in randomized order. Mice were allowed to freely bite the flavored food or sponge pieces during 5-minute sessions while calcium activity was continuously recorded.

### Linguine feeding assay with optogenetic manipulation

Mice were fasted for 4 hours prior to testing to enhance motivation. During each session, animals were individually placed in a clean behavioral arena and presented with three pre-weighed pieces of raw linguine (approximately 2 ⍰ cm each). Mice were allowed to freely interact with and consume the pasta for 10 minutes. Each mouse underwent two test sessions on separate days in a counterbalanced order: one session with light stimulation and one without. For the optogenetic activation experiment, 5 Hz, 10 H and 20 Hz blue light (473 ⍰ nm) was delivered throughout the 10-minute session. For the inhibition experiment, yellow light (532 ⍰ nm) was delivered continuously throughout the 10-minute test. Residual linguine was weighed after each session to determine food consumption.

### Biting behavior assay with food or Styrofoam

Mice were presented with small objects made of either regular chow or styrofoam. Prior to the test, five uniform cubes (approximately 0.5 ⍰ cm × 0.5 ⍰ cm × 0.5 ⍰ cm) of food or styrofoam were placed in a clean transparent chamber. Mice were allowed to freely interact with the objects for the duration of the session. Optogenetic stimulation was delivered in a structured alternating pattern of 1 ⍰ minute OFF followed by 1 minute ON, repeated three times over a total of 6 minutes. Blue light (473 ⍰ nm) was delivered via a fiber-optic patch cable connected to a laser through a rotary joint, with light power calibrated at the fiber tip (10–15 ⍰ mW). Light delivery was controlled using a pulse generator to achieve consistent timing across animals. Bottom-up video capture of the mouse’s behavior via a camera positioned below the chamber and biting events were scored manually.

### Cell registration

To identify the same individual cells across diverse imaging settings, we used centroid extraction, stiff alignment, and optimal one-to-one matching. For each imaging condition, we first extracted spatial footprints from the binarized cell maps and computed the centroid of each segmented footprint. One condition was treated as the reference map, and centroids from the other condition were co-registered to this reference using a rigid transformation (translation and rotation) estimated from the top nearest-neighbor centroid pairs via singular value decomposition. Following alignment, we computed the whole pairwise centroid-distance matrix between conditions and used the Hungarian approach to achieve the best one-to-one assignment; cell pairings with a post-registration centroid distance of less than 35 pixels were considered matched.

### PCA

We performed principal component analysis (PCA) on event-aligned population activity to characterize the dominant low-dimensional structure. For each neuron, z-scored calcium traces were extracted from –4 to +6 s relative to bite onset. Neurons with sparse or unreliable responses were excluded by requiring that > 5 % of samples exceeded zero across trials. Activity in each condition was baseline-aligned by subtracting the mean signal from a pre-event window (– 0.5 to 0 s), and peak-normalized across conditions so that no single unit dominated the population embedding. To estimate the shared low-dimensional axes, we constructed a neuron × time matrix by concatenating post-bite activity (0 to + 5 s) across conditions and centered each time bin before decomposition. Condition-specific trajectories were then obtained by projecting the event-aligned activity back onto the PCA loadings. Importantly, no temporal smoothing was applied prior to PCA; traces were only spline-interpolated for visualization, and all loadings and variance estimates were computed from unsmoothed signals.

### Pairwise Pearson correlation analysis

To quantify representational similarity across conditions, we computed Pearson correlation coefficients based on condition-averaged neural activity. For each neuron, we extracted z-scored calcium signals within a – 6 s to + 8 s event window. Mean z-scored activity was calculated for each neuron × condition, yielding a neuron-by-condition response matrix. Neural responses were averaged across neurons for each material condition, and pairwise Pearson correlations were computed between material-level response vectors to quantify similarity across conditions.

### Statistical analysis

All statistics are described where used. Statistical analyses were conducted using GraphPad Prism 9 software (GraphPad). No statistical methods were used to predetermine sample sizes, but the number of samples in each group were similar to those reported in previous publications. Data collection and analysis were not performed blind to the conditions of the experiments. Mice that, after histological inspection, had the location of the viral injection (reporter protein) or of the optic fiber (s) outside the area of interest were excluded. All data were represented as the mean ± SEM. Significance levels are indicated as follows: *p < 0.05; **p < 0.01; ***p < 0.001; ****p < 0.0001.

---

**Figure. S1.**
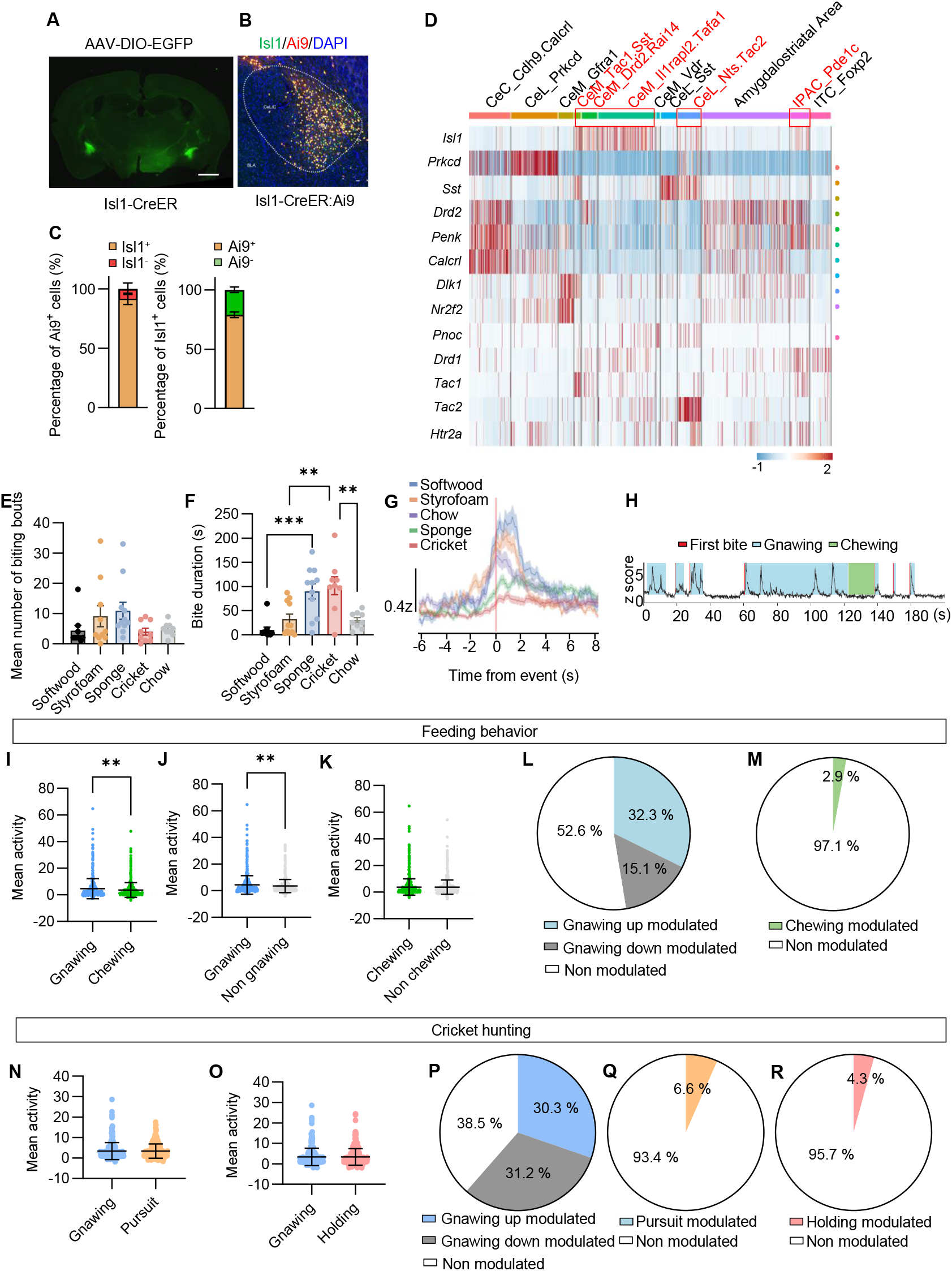
CeA^Isl1^ neurons were activated at the bite onset. **A**. Representative histological image of EGFP expression in the CeA. Scale bar, 100 µm. **B**. Representative image showing Isl1 immunostaining (green) and Ai9 reporter expression (red) with DAPI (blue) in an Isl1-CreER: Ai9 mouse. The CeA subregions are outlined. Scale bar, 30 µm. **C**. Quantification of the proportion of Ai9^+^ neurons expressing Isl1 and the proportion of Isl1^+^ neurons expressing Ai9 (n = 3 mice). **D**, Heatmap of marker gene expression for CeA neuronal subpopulations. Cells are grouped and color-coded by transcriptionally defined clusters (top), as described in a previous study^1^. Red outlines and red text highlight the Isl1-positive population. **E**. Quantification of mean number of biting bouts for objects with varying physical properties (n=10 mice; N.S.; one-way ANOVA test) **F**. Quantification of bite duration for objects with varying physical properties (n=10 mice; ***P <0.001, **P <0.01; one-way ANOVA test). **G**. Average responses of the z-scored calcium activity for each condition: softwood, Styrofoam, sponge, cricket, and chow. Shaded areas represent s.e.m. **H**. Representative z-scored calcium trace from a single neuron aligned to behavioral events: first bite (red), eating (blue), and chewing (green). **I-M**, Feeding behavior. **I**. Mean neuronal activity during gnawing and chewing epochs (n=10 mice; **P < 0.01; Wilcoxon signed-rank test). **J**. Mean neuronal activity during gnawing versus non-gnawing epochs (n=10 mice; **P < 0.01; Wilcoxon signed-rank test). **K**. Mean neuronal activity during chewing versus non-chewing epochs (n=10 mice; N.S.; Wilcoxon signed-rank test). **L**. Proportion of neurons up-modulated, down-modulated, and non-modulated during gnawing. **M**. Proportion of neurons modulated during chewing. **N-R**. Cricket hunting behavior. **N**. Mean neuronal activity during gnawing and pursuit epochs (n=10 mice; N.S.; Wilcoxon signed-rank test). **O**. Mean neuronal activity during gnawing and holding epochs (n=10 mice; N.S.; Wilcoxon signed-rank test). **P**. Proportion of neurons up-modulated, down-modulated, and non-modulated during gnawing. **Q**. Proportion of neurons modulated during pursuit. **R**. Proportion of neurons modulated during holding.

**Figure. S2.**
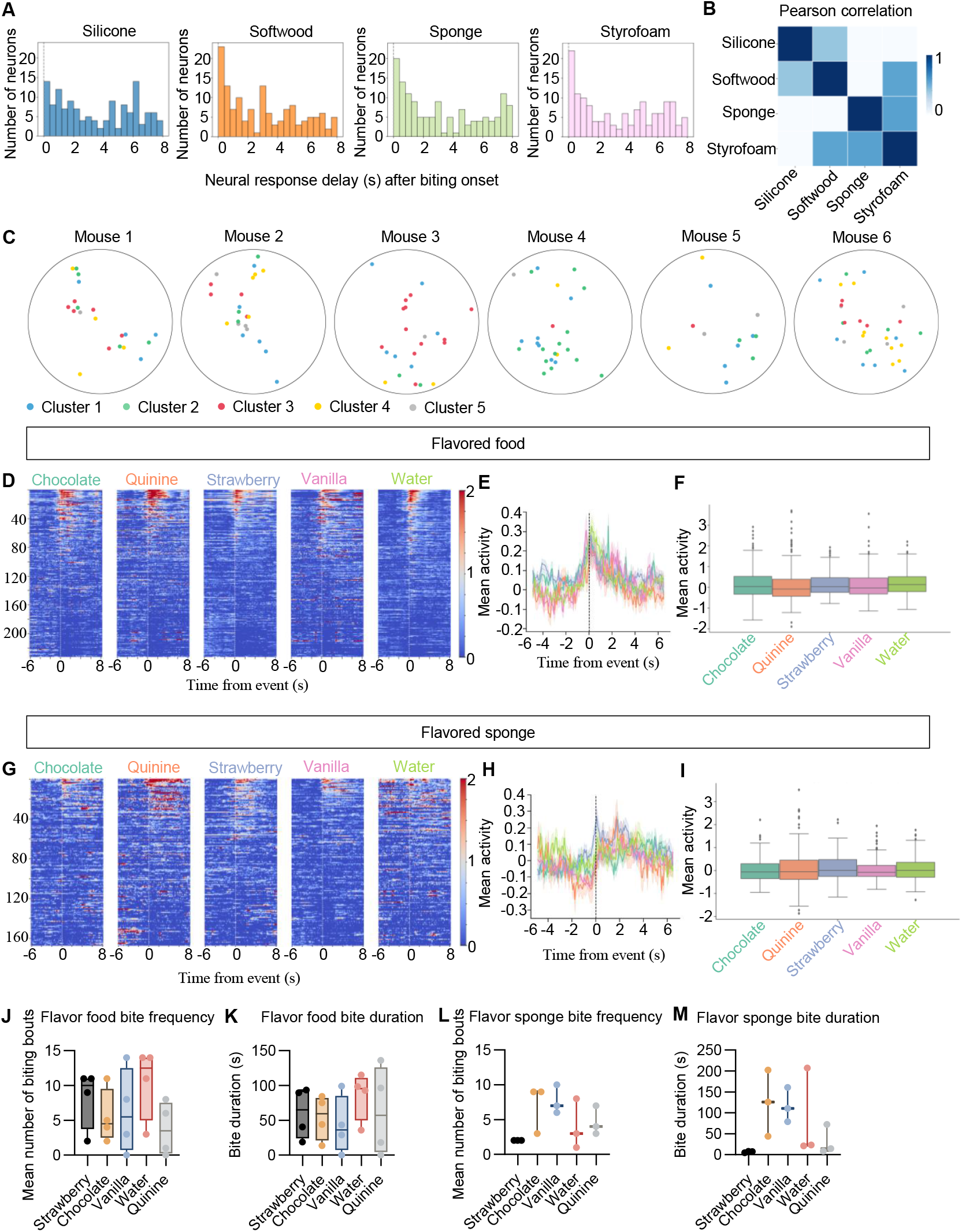
Neuronal responses to flavored food and flavored sponge. **A**. Histograms showing the distribution of response delays (s) after biting onset for neurons recorded during interactions with silicone (blue), softwood (orange), sponge (green), and Styrofoam (purple). Each bar indicates the number of neurons exhibiting peak activity within the corresponding time bin. **B**. Correlation coefficient matrixes of the responses of all neurons for each condition. **C**. The spatial locations of individual extracted neurons in the field of view (FOV) in the CeA of six mice. clusters labeled by different color code. **D**. Heatmaps showing normalized activity of individual neurons aligned to bite onset for different flavored food. Each row represents one neuron, and color scale indicates z-scored activity (n= 4 mice). **E**. Mean population activity traces for each flavored food, showing the average z-scored activity aligned to bite events. Shaded areas represent ± s.e.m. **F**. Box plots summarizing mean activity across different flavored food (n=4 mice; N.S.; Wilcoxon signed-rank test). **G**. Heatmaps showing normalized activity of individual neurons aligned to bite onset for different flavored sponge. Each row represents one neuron, and color scale indicates z-scored activity (n= 3 mice). **H**. Mean population activity traces for each flavored sponge, showing the average z-scored activity aligned to bite events. Shaded areas represent ± s.e.m. **I**. Box plots summarizing mean activity across different flavored sponge (n=3 mice; N.S.; Wilcoxon signed-rank test). **J**. Quantification of mean number of biting bouts for different flavored food (n=4 mice; N.S.; Wilcoxon signed-rank test). **K**. Quantification of bite duration for different flavored food (n=4 mice; N.S.; Wilcoxon signed-rank test). **L**. Quantification of mean number of biting bouts for different flavored sponge (n=3 mice; N.S.; Wilcoxon signed-rank test). **M**. Quantification of bite duration for different flavored sponge (n=3 mice; N.S.; Wilcoxon signed-rank test).

**Figure. S3.**
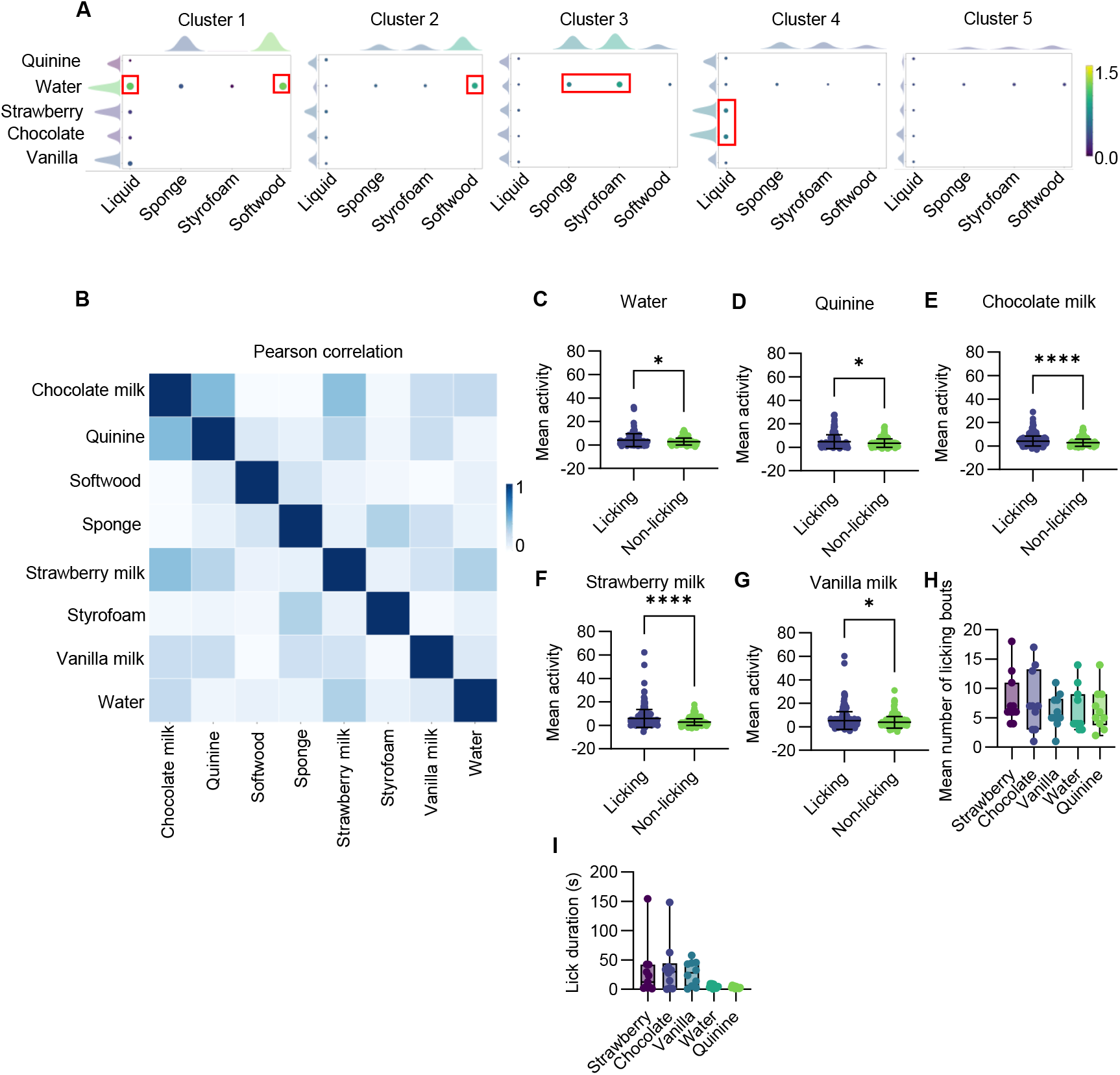
The encoding properties of CeA^Isl1^ neurons. **A**. Summary matrix of mean peak response across stimuli in each cluster. Dot size and color represent average response magnitude, and density plots indicate the distribution of neuron responses to each stimulus. **B**. orrelation coefficient matrixes of the responses of all neurons for each condition. **C-G**. Mean neuronal activity during licking versus non-licking epochs: water (**C**), quinine (**D**), chocolate milk (**E**), strawberry milk (**F**), and vanilla milk (**G**) (n= 6 mice; *P < 0.05, ****P < 0.0001, Wilcoxon signed-rank test). **H**. Quantification of mean number of licking bouts or different flavored milk (n=6 mice; N.S.; one-way ANOVA test). **I**. Quantification of lick duration for different flavored milk (n=6 mice; N.S.; one-way ANOVA test).

**Figure. S4.**
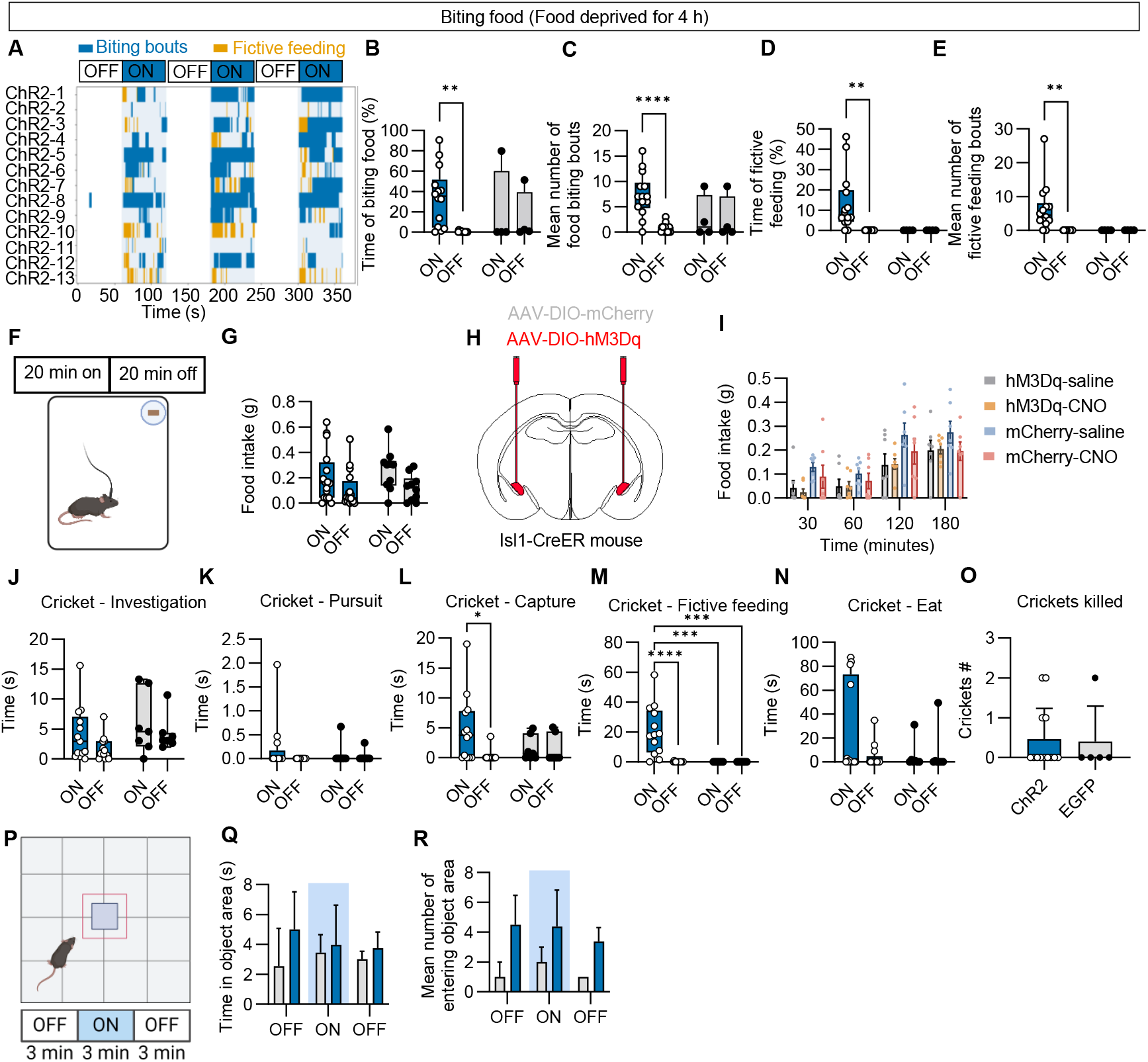
Activation of CeA^Isl1^ neurons in feeding, hunting, and exploratory behaviors. **A**. Raster plot showing biting bouts (blue) and fictive feeding (orange) across individual 4-h food-deprived ChR2 mice during alternating light OFF and light ON epochs in biting food assay. **B-E**. Quantification of behaviors during light ON and OFF periods for both ChR2 mice (n = 14) and EGFP mice (n = 4) in food biting assay. **B**. Percentage of time spent biting food. **C**. Mean number of f food-biting bouts. **D**. Percentage of time engaged in fictive feeding. **E**. Mean number of fictive feeding bouts. Two-way ANOVA with Tukey’s post hoc test, **P < 0.01, ****P < 0.0001. Blue is ChR2 group, and grey is EGFP. **F**. Schematic of the paradigm for testing the effects of photoactivation on feeding behavior. **G**. Food intake by fed ChR2 mice (n = 16) and EGFP mice (n = 9). N.S.; two-way ANOVA with Tukey’s post hoc test. **H**. Schematic of CeA^Isl1^ neurons expressing either the excitatory DREADD hM3Dq or control mCherry. **I**. Cumulative food intake of fed Isl1-CreER animals that expressing the excitatory DREADD hM3Dq or control mCherry in the CeA, after i.p. injections of saline and CNO (2 mg/Kg). N = 7 mice per group; N.S.; two-way ANOVA with Tukey’s post hoc test. **J-O**. Quantification of different behavioral epochs during light ON and OFF period for both ChR2 mice (n = 13) and EGFP mice (n = 7) in cricket hunting tasks. **J**. Time spent investigating crickets. **K**. Time spent pursuing crickets. **L**. Time spent capturing crickets. **M**. Time spent on fictive feeding. **N**. Time spent eating crickets. **O**. Number of crickets killed. Two-way ANOVA with Tukey’s post hoc test, *P < 0.05, ***P < 0.0001, ****P < 0.0001. **P**. Schematic of the paradigm for testing the effects of photoactivation on novel object exploration. **Q**. Time spent in the object area for both ChR2 mice (n = 8) and EGFP mice (n = 3); N.S.; two-way ANOVA with Tukey’s post hoc test. **R**. Mean number f entering the object area for both ChR2 mice (n = 8) and EGFP mice (n = 3) N.S.; two-way ANOVA with Tukey’s post hoc test.

**Figure. S5.**
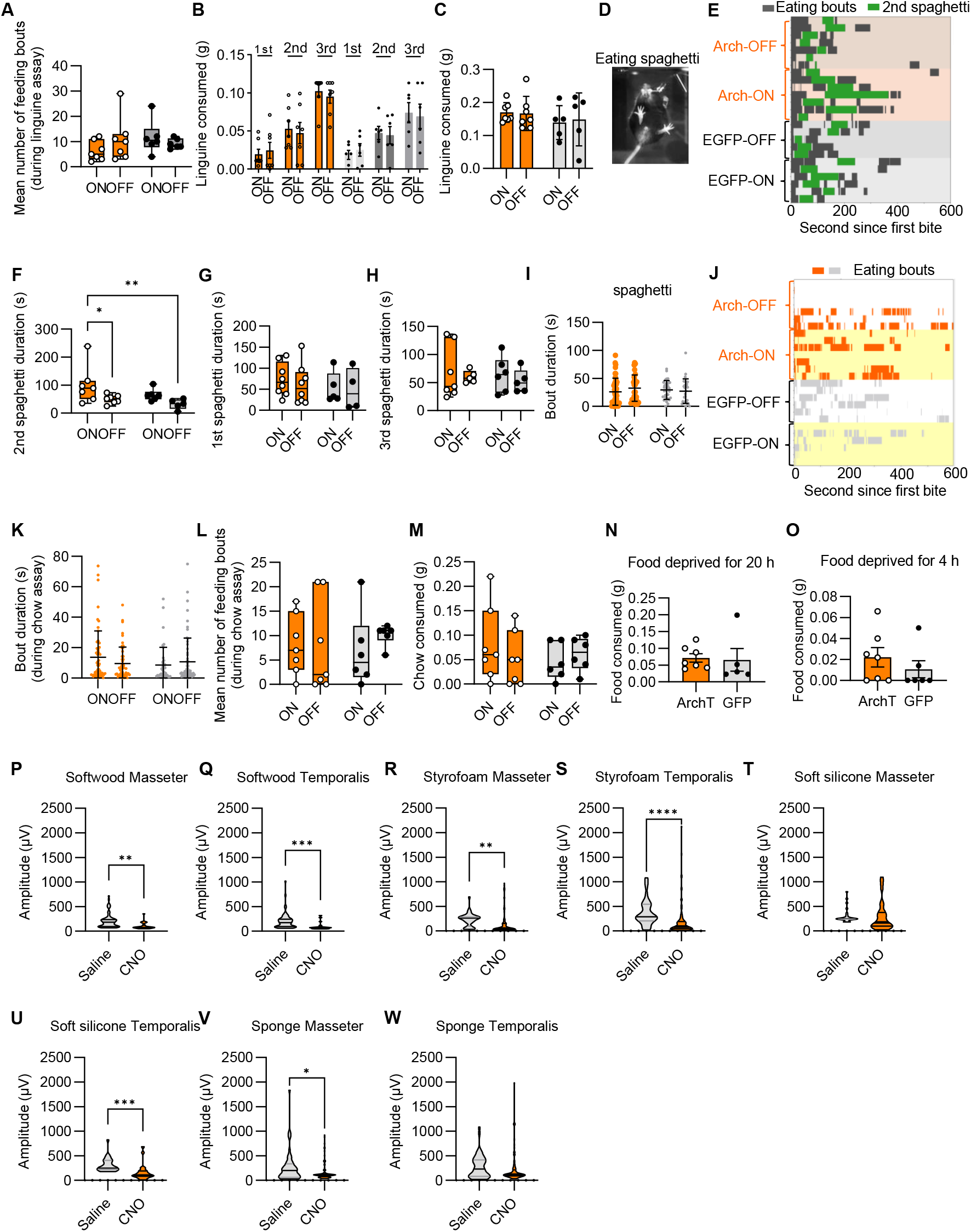
Inhibition of CeA^Isl1^ neurons does not affect food consumption. **A**. Mean number of feeding bouts across light OFF and ON epochs in Arch3.0 and EGFP groups during linguine consumption assays (N.S.; two-way ANOVA with Tukey’s post hoc test). **B**. Cumulative consumption of the first, second, and third linguine pieces across light OFF and ON epochs in Arch3.0 and EGFP groups (N.S.; two-way ANOVA with Tukey’s post hoc test). **C**. Linguine intake by 4-h food-deprived Arch3.0 mice (n = 7) and EGFP mice (n = 5) during alternating light OFF and light ON epochs. N.S.; two-way ANOVA with Tukey’s post hoc test. **D**. Example frame of a mouse consuming spaghetti. **E**. Raster plots showing eating bouts (black) and consumption duration of the second spaghetti piece (green) across light OFF and ON epochs in Arch3.0 and EGFP groups during spaghetti consumption assays. **F**. Duration of second spaghetti consumption across light OFF and ON epochs in Arch3.0 and EGFP groups (n = 7 mice per group; *P < 0.05, **P < 0.01; Two-way ANOVA with Tukey’s post hoc test). **G**. Duration of first spaghetti consumption across light OFF and ON epochs in Arch3.0 (n = 8) and EGFP (n = 5) groups. N.S.; two-way ANOVA with Tukey’s post hoc test. **H**. Duration of third spaghetti consumption across light OFF and ON epochs in Arch3.0 (n = 8) and EGFP (n = 5) groups. N.S.; two-way ANOVA with Tukey’s post hoc test. **I**. Duration of individual feeding bouts across light OFF and ON epochs in Arch3.0 (n = 8) and EGFP (n = 5) groups during spaghetti consumption assays. N.S.; two-way ANOVA with Tukey’s post hoc test. **J**. Raster plots showing eating bouts (orange for Arch3.0 and grey for EGFP) across light OFF and ON epochs in Arch3.0 and EGFP groups during chow consumption assays. **K**. Duration of individual feeding bouts across light OFF and ON epochs in Arch3.0 and EGFP groups during chow consumption assays (n = 7 mice per group; N.S.; Two-way ANOVA with Tukey’s post hoc test). **L**. Mean number of feeding bouts across light OFF and ON epochs in Arch3.0 and EGFP groups during chow consumption assays (n = 7 mice per group; N.S.; Two-way ANOVA with Tukey’s post hoc test). **M**. Cumulative consumption of the chow across light OFF and ON epochs in Arch3.0 and EGFP groups (n = 7 mice per group; N.S.; Two-way ANOVA with Tukey’s post hoc test). **N**. Food intake by 20-h food-deprived Arch3.0 mice (n = 7) and EGFP mice (n = 5). N.S.; two-way ANOVA with Tukey’s post hoc test. **O**. Food intake by 4-h food-deprived Arch3.0 mice (n = 7) and EGFP mice (n = 5). N.S.; two-way ANOVA with Tukey’s post hoc test. **P**. EMG amplitude measurements from the masseter muscle during softwood biting after saline or CNO (0.4 mg kg^−1^) injection in Isl1-CreER; hM4Di mice. n=4; **P < 0.01; paired t-test. **Q**. EMG amplitude measurements from the temporalis muscle during softwood biting after saline or CNO (0.4 mg kg^−1^) injection in Isl1-CreER; hM4Di mice. n=4; ***P < 0.001; paired t-test. **R**. EMG amplitude measurements from the masseter muscle during Styrofoam biting after saline or CNO (0.4 mg kg^−1^) injection in Isl1-CreER; hM4Di mice. n=4; **P < 0.01; paired t-test. **S**. EMG amplitude measurements from the temporalis muscle during Styrofoam biting after saline or CNO (0.4 mg kg^−1^) injection in Isl1-CreER; hM4Di mice. n=4; ****P < 0.0001; paired t-test. **T**. EMG amplitude measurements from the masseter muscle during soft silicone biting after saline or CNO (0.4 mg kg^−1^) injection in Isl1-CreER; hM4Di mice. n=4; N.S; paired t-test. **U**. EMG amplitude measurements from the temporalis muscle during soft silicone biting after saline or CNO (0.4 mg kg^−1^) injection in Isl1-CreER; hM4Di mice. n=4; ***P < 0.001; paired t-test. **V**. EMG amplitude measurements from the masseter muscle during sponge biting after saline or CNO (0.4 mg kg^−1^) injection in Isl1-CreER; hM4Di mice. n=4; *P < 0.05; paired t-test. **W**. EMG amplitude measurements from the temporalis muscle during sponge biting after saline or CNO (0.4 mg kg^−1^) injection in Isl1-CreER; hM4Di mice. n=4; N.S; paired t-test.

**Figure. S6.**
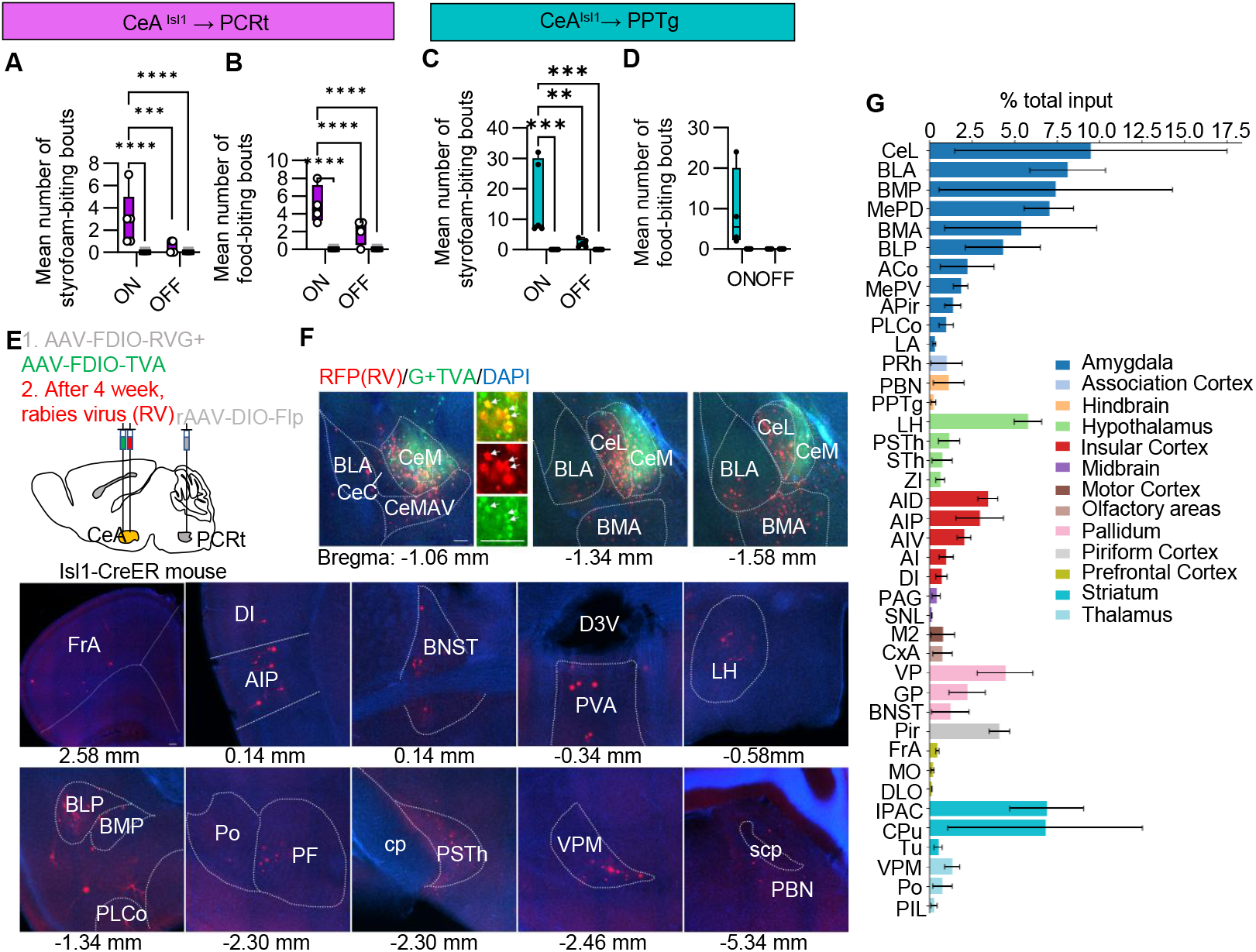
Brain network architecture of CeA^Isl1^ neurons. **A**. Mean number of Styrofoam-biting bouts during ON and OFF periods for both CeA ^Isl1^ → PCRt ChR2 (pink) mice and EGFP (grey) mice in biting assay. N = 5 mice per group. Two-way ANOVA with Tukey’s post hoc test, ***P < 0.001, ****P < 0.0001. **B**. Mean number of food-biting bouts during ON and OFF periods for both CeA ^Isl1^ → PCRt ChR2 (pink) mice and EGFP (grey) mice in biting assay. N = 5 mice per group. Two-way ANOVA with Tukey’s post hoc test, ****P < 0.0001. **C**. Mean number of Styrofoam-biting bouts during ON and OFF periods for both CeA ^Isl1^ → PPtg ChR2 (green) mice and EGFP (grey) mice in biting assay. N = 5 mice per group. Two-way ANOVA with Tukey’s post hoc test, **P < 0.01, ***P < 0.001. **D**. Mean number of food-biting bouts during ON and OFF periods for both CeA ^Isl1^ → PPtg ChR2 (green) mice and EGFP (grey) mice in biting assay. N = 5 mice per group. Two-way ANOVA with Tukey’s post hoc test, N.S. **E**. Schematic of the strategy for monosynaptic retrograde rabies virus (RV) tracing. **F**. Representative image of the injection site and input areas. High-magnification panel indicate starter cells expressing G + TVA (green) and RV (red) (arrows). Scale, 30 µm. **G**. Major brain regions projecting to CeA^Isl1^ neurons depicted as percentage of total inputs (n = 3 mice). **Abbreviations:** ACo, anterior cortical amygdaloid nucleus; AI, agranular insular cortex; AID, agranular insular cortex, dorsal part; AIP, agranular insular cortex, posterior part; AIV, agranular insular cortex, ventral part; APir, amygdalopiriform transition area; BNST, the bed nucleus of the stria terminalis; BLA, basolateral amygdaloid nucleus, anterior part; BLP, basolateral amygdaloid nucleus, posterior part; BMP, basomedial amygdaloid nucleus, posterior part; BMA, basomedial amygdaloid nucleus, anterior part; CeL, lateral part of central amygdala; CPu, caudate putamen (striatum); CxA, cortex-amygdala transition zone; DI, dysgranular insular cortex; DLO, dorsal lateral olfactory tract; D3V, dorsal 3rd ventricle; FrA, frontal association cortex; GP, globus pallidus; IPAC, interstitial nucleus of the posterior limb of the anterior commissure; LA, lateral amygdaloid nucleus; LH, lateral hypothalamic area; MePD, medial amygdaloid nucleus, posterodorsal part; MePV, medial amygdaloid nucleus, posteroventral part; M2, secondary motor cortex; MO, medial orbital cortex; PAG, periaqueductal gray; PBN, parabrachial nucleus; PPTg, pedunculopontine tegmental nucleus; PLCo, posterolateral cortical amygdaloid nucleus; PVA, paraventricular thalamic nucleus, anterior part; Po, posterior thalamic nuclear group; PIL, posterior intralaminar thalamic nucleus; PRh, perirhinal cortex; PSTh, parasubthalamic nucleus; Pir, piriform cortex; SNL, substantia nigra, lateral part; STh, subthalamic nucleus; scp superior cerebellar peduncle (brachium conjunctivum); Tu, olfactory tubercle; VP, ventral pallidum; VPM, ventral posteromedial thalamic nucleus; ZI, zona incerta.

**Figure. S7.**
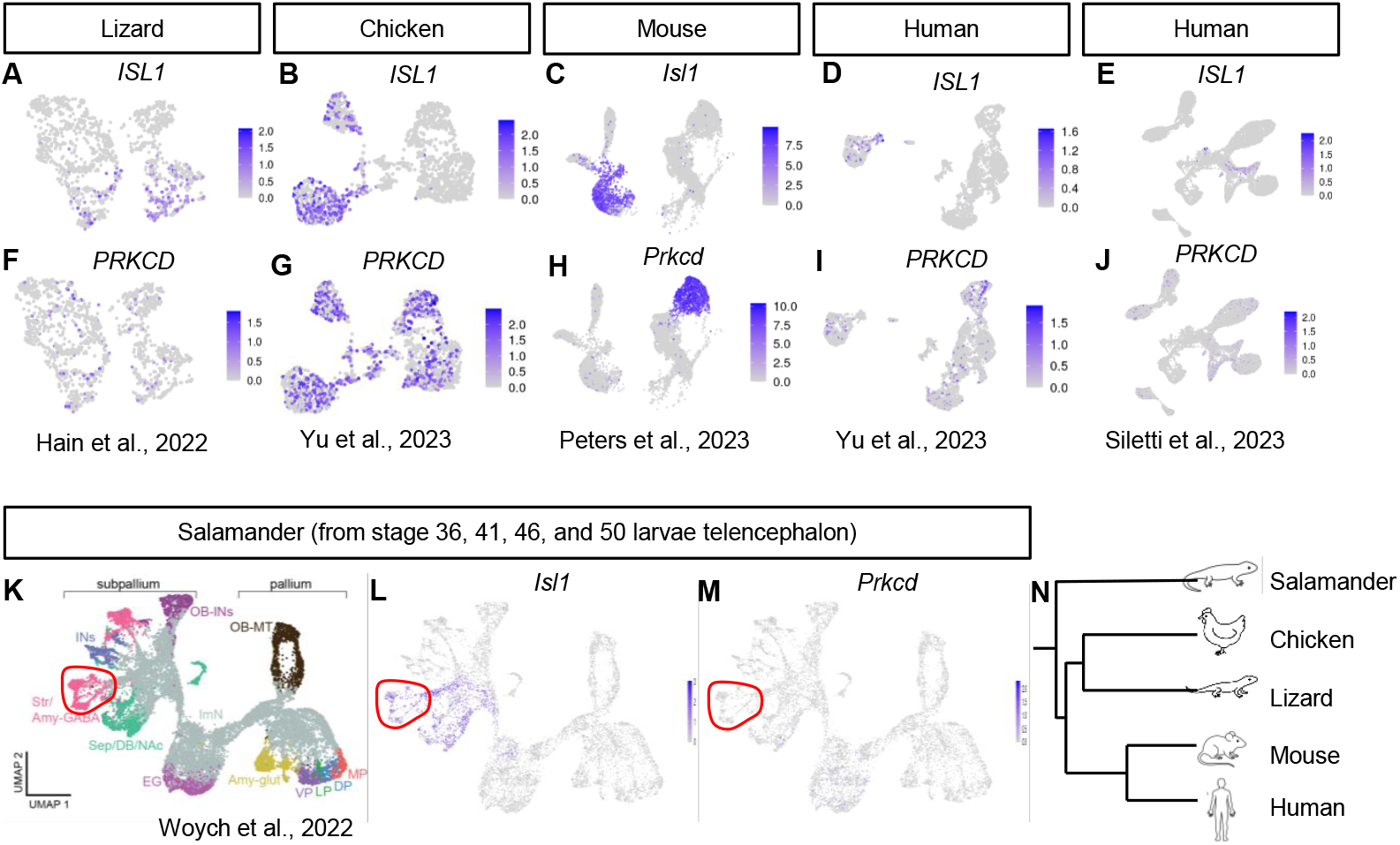
Conserved segregation of *Isl1* and *Prkcd* neuronal populations in the central amygdala across vertebrates. **A–J**, Single-cell transcriptomic maps showing expression of *Isl1* (**A-E**) and *Prkcd* (**F-J**) across CeA clusters in lizard^2^, chicken^3^, mouse^1^, and human^3,4^. **K–M**, UMAP visualization of salamander telencephalon (stages 36, 41, 46, and 50)^5^ showing annotated brain regions (**K**) and expression of *Isl1* (**L**) and *Prkcd* (**M**). Red outlines indicate putative CeA regions. **N**, Schematic phylogenetic tree illustrating the species analyzed.

